# The PD-1 checkpoint receptor maintains tolerance of self-reactive CD8 T cell in skin

**DOI:** 10.1101/2021.07.09.451765

**Authors:** Martina Damo, Can Cui, Ivana William, Noah I. Hornick, Darwin Kwok, Kathryn Clulo, William E. Damsky, Jonathan S. Leventhal, Nikhil S. Joshi

**Affiliations:** Department of Immunobiology, Yale University School of Medicine, New Haven, CT 06519, USA; Department of Dermatology, Yale University School of Medicine, New Haven, CT 06519, USA; Department of Pathology, Yale University School of Medicine, New Haven, CT 06519, USA

**Keywords:** Tolerance, immune checkpoint receptors, PD-1, CD8 T cells, irAEs

## Abstract

Peripheral tolerance is thought to result from anergy or deletion of self-reactive T cells shortly after antigen encounter. However, the frequent occurrences of immune-related Adverse Events (irAEs) following checkpoint inhibitor (CPI) treatment suggest a hypothesis that immunologically healthy individuals have self-reactive effector T cells that are kept in a non-pathogenic state through checkpoint receptor-mediated suppression, instead of anergy or deletion. We expressed self-antigens in healthy skin and found that antigen-specific CD8 T cells infiltrated the tissue, but remained tolerant, despite having a transcriptional program that resembled effector T cells found after CPIs. These self-reactive PD-1+ CD8 T cells drove IFNγ-dependent increases in PD-L1 on skin myeloid cells. Blockade of PD-1 or PD-1/CTLA-4 led to post-transcriptional upregulation of effector proteins by antigen-specific CD8 T cells and elimination of antigen-expressing epithelial cells, resulting in localized tissue pathology with features of human cutaneous irAEs. This data supports the hypothesis that myeloid cells in healthy skin prevent pathology from self-reactive effector CD8 T cells through the PD-1/PD-L1 pathway.

## Introduction

The widespread use of immune checkpoint inhibitors (CPIs) in cancer patients has been a revolution for oncology (Sharma and Allison). CPIs are now used as a standard therapy across many cancer types, but their widespread use has also led to a sharp rise in immune-related Adverse Events (irAEs). irAEs can occur in up to 76% of CPI-treated patients and affect many organ systems throughout the body (Dougan et al.; Postow et al.; Ramos-Casals et al.). Yet, little is known about the mechanisms causing irAEs. A broad range of effector molecules and cell types could be involved in irAEs, but many are likely T cell driven. This includes cutaneous lichenoid irAEs, colitis, myocarditis, and thyroiditis, which have clear T cell involvement, based on significant T cell accumulation, and even clonal T cell expansion, in affected tissues (Coleman et al.; Johnson et al.; Luoma et al.; Shi et al.; Yasuda et al.). Moreover, pathogenic self-reactive T cells could be driving some irAEs. Most irAEs resemble localized autoimmune diseases, and a handful of rare irAEs share genetic risk alleles with their autoimmune counterparts (Khan et al.; Stamatouli et al.; van Belle et al.). However, the fact that most CPI-treated patients have some form of irAE suggests that many irAEs may not result from simple exacerbation of pre-existing, but sub-clinical, autoimmune disease (Luoma *et al.*; Postow *et al.*; Smillie et al.). This raises the intriguing hypothesis that immune checkpoint receptors (ICRs) like PD-1 and CTLA-4 may have an active role in maintaining peripheral tolerance of self-reactive T cells in non-autoimmune prone individuals.

Most self-reactive CD8 or CD4 T cells are eliminated by negative selection during thymocyte development (central tolerance; Klein et al.). However, CD8 T cells with self-antigen (Ag)-specific T cell receptors (TCRs) are present in healthy people, and these cells exist at similar frequencies to CD8 T cells that are specific for foreign Ags (Maeda et al.; Yu et al.). Despite their presence, self-reactive T cells do cause immunopathology in most people. This is due to the existence of peripheral tolerance mechanisms. Peripheral T cell tolerance is mediated by both T cell-intrinsic (anergy and clonal deletion) and T cell-extrinsic (regulatory T cells, Tregs) mechanisms (ElTanbouly and Noelle). T cell-intrinsic mechanisms result from self Ag presentation by tolerogenic dendritic cells (DCs), which fail to provide enough antigen and costimulatory signals to generate effector T cells (Probst et al.; Redmond et al.; Redmond and Sherman). Intrinsic-tolerized self-reactive T cells do not acquire effector functions and often cannot enter the self-antigen expressing tissue (Bianchi et al.; Parish et al.). Seminal work showed ICR-deficiency or blockade could prevent the establishment of T cell-intrinsic tolerance for self-reactive CD8 T cells in well-characterized animal models of T-cell anergy and deletional tolerance (Keir et al.; Keir et al.; Kurts et al.; Martin-Orozco et al.; Nelson et al.). However, CPIs also did not rescue self-reactive T cells after intrinsic tolerance was established, either because the self-reactive T cells were unresponsive to treatment (in the case of anergy) or because they no longer existed (in the case of deletion). Likewise, extensive analysis of ICR-deficient mice (*i.e., Pdcd1−/−* mice) showed the development of spontaneous autoimmune diseases, particularly when mice were on autoimmune prone backgrounds, suggesting that interactions between ICR-expressing T cells and their ligands could play a T cell-extrinsic role in delaying or dampening ongoing autoimmune disease processes (Tivol et al.; Waterhouse et al.; Brown et al.; Keir *et al.*; Keir *et al.*; Latchman et al.; Martin-Orozco *et al.*; Nishimura et al.; Nishimura et al.; Tivol *et al.*; Wang et al.). Here, while T cells express ICRs, this mechanism is “extrinsic” because ICRs do not provide T cells with ligand-independent signals (they act by ligand-dependent dampening positive signals T cells see from Ag/costimulatory receptors). ICRs are also not expressed until after T cell activation. However, in ICR-deficient models, it was difficult to dissociate the impacts of peripheral and central T cell tolerance, and thus it remained uncertain if their spontaneous disease was due to defects in central tolerance that allowed self-reactive T cells to enter the peripheral naïve T cell pool, defects in establishing and/or maintaining peripheral tolerance, or all of the above. By contrast, CPI treatment of wild-type mice was extensively studied in preclinical tumor models, and autoimmune-like side effects were rarely observed (if ever) (Grosso and Jure-Kunkel; Okazaki and Honjo). Thus, the frequent appearance of irAEs in CPI-treated patients was surprising, as it suggested that substantial fractions of patients had T cells that were capable of causing pathology, but that this pathology was being held in check by the functions of the ICRs and their ligands.

Given the complications with existing peripheral tolerance models, most models for irAEs have involved CPI treatment of autoimmune-prone or disease-induced mice, which accelerated disease onset or exacerbated disease severity and was useful for testing therapies to ameliorate irAEs (Perez-Ruiz et al.; Yasuda *et al.*). However, because disease was observed regardless of CPI treatment, these models could not inform on the mechanisms for how CPIs broke peripheral tolerance. Thus, to investigate potential irAE-causing mechanisms, we developed a novel animal model for inducing peripheral T cell tolerance based on the recently published iNversion INducible Joined neoAntigen (NINJA) mouse model (Damo et al.). NINJA allows for *de novo* induction of self Ag expression in a peripheral tissue, without the confounding influence of central tolerance on the endogenous naïve NINJA-Ag-specific CD4 or CD8 T cells. Critically, NINJA mice are on a non-autoimmune prone C57BL/6 (B6) background, and thus the expectation was that self Ag induction in peripheral organs would result in peripheral tolerance of endogenous Ag-specific T cells, allowing for deeper mechanistic investigations into how, when, and where tolerance occurs. Here, we use induction of self-Ag in skin to study the mechanisms mediating peripheral T cell tolerance and the role of CPIs in development of irAEs.

## Results

### Ag expression in skin results in accumulation of Ag-specific CD8 T cells and tolerance

To study the impact of *de novo* expression of self Ag in skin, we bred NINJA x CAG-rtTA3 mice (henceforth referred to as NINJA) to mice expressing Cre recombinase fused to the estrogen receptor (CreER^T2^) from the universally expressed endogenous promoter in the *Rosa26* locus to generate “N/C” mice (*Rosa26-NINJA/CreER^T2^;CAG-rtTA3 Tg*). Administration of doxycycline (Dox; food) and 4-hydroxy-tamoxifen (Tam; applied topically on a selected area of the skin) to N/C mice induced a series of genetic recombination events and resulted in the expression of genetically encoded Ag (LCMV-derived GP_33-43_ and GP_61-80_) embedded in a molecule of GFP, which was localized to the Tam-treated skin area (**Fig 1A**). We previously showed that untreated N/C mice were not tolerant to the Ag in NINJA because the naïve endogenous GP33-specific CD8 and GP66-specific CD4 T cells in N/C mice responded similarly to their counterparts in B6 mice after acute infection with LCMV (Damo *et al.*). Dox/Tam treatment of N/C mice led to responses by endogenous GP33-specific CD8 T cells in the Tam-treated skin and local draining lymph nodes (dLNs) at day 15-16 post Dox/Tam (**Fig 2A**) and skin-infiltrating GP33-specific CD8 T cells were still detectable in skin 30+ days after Ag induction (not shown). Yet, despite their infiltration into skin, the macroscopic appearance of Ag-expressing skin in Dox/Tam-treated N/C mice (+ Ag) was comparable to skin in “No Ag” negative controls (**Fig 1C**; note, as No Ag controls, we used Dox/Tam-treated B6 or NINJA mice and untreated N/C mice. These were phenotypically identical to each other). Together, these data showed that self Ag-specific T cells in the N/C model were not deleted, but rather they infiltrated the Ag-expressing tissue. However, despite this infiltration, local tissue homeostasis and immunological tolerance were maintained.

**Figure 1.**
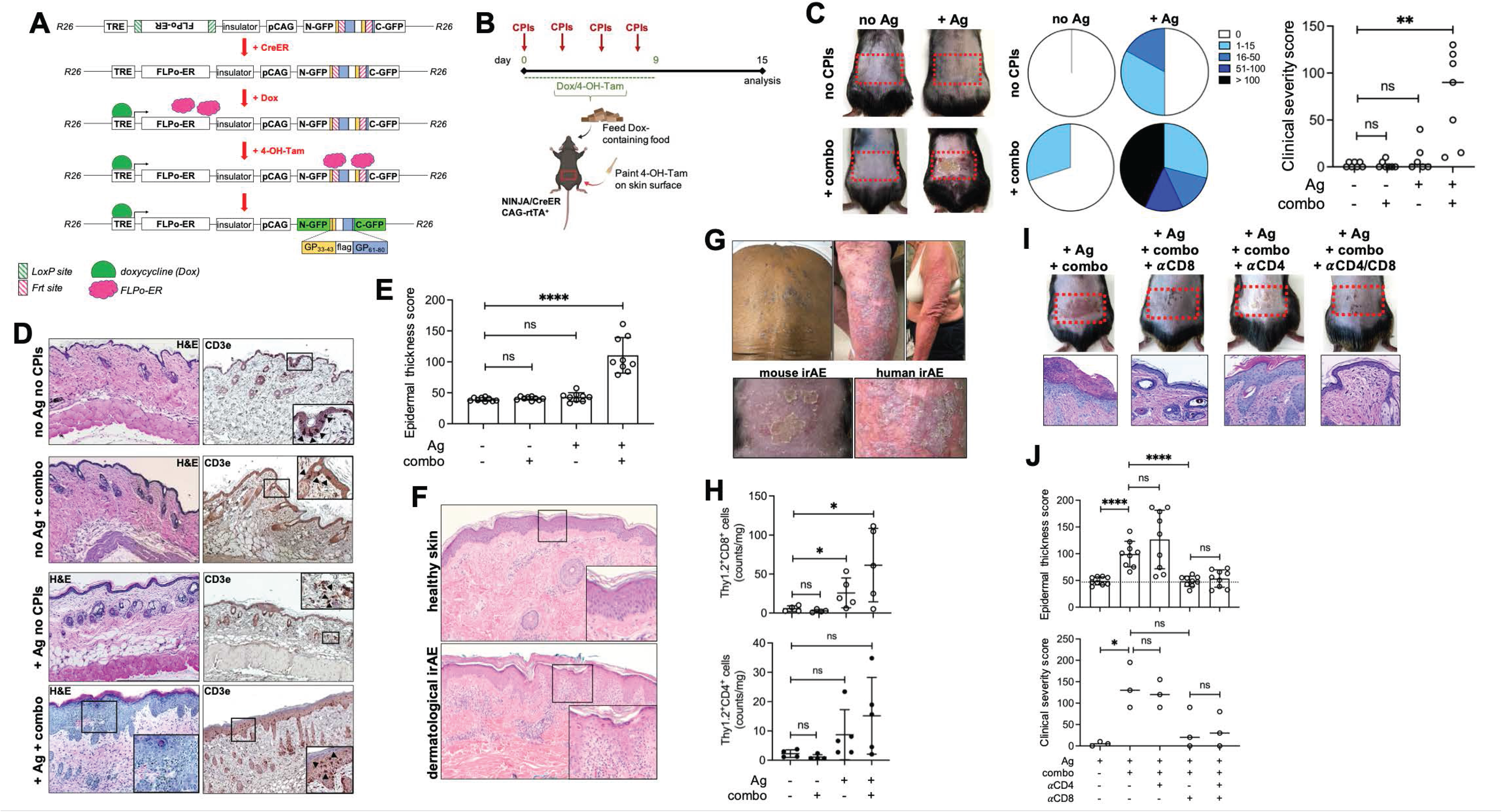
Skin-specific Ag expression coupled with CPI administration leads to CD8 T cell-dependent local lichenoid reaction-like tissue pathology. A) Ag expression in the NINJA x CAG-rtTA3 (henceforth referred to as NINJA) mouse model is dependent on Cre or CreER activity and administration of doxycycline (Dox) and 4-hydroxy-Tamoxifen (Tam). B) Experimental schedule: NINJA x R26-CreER^T2^ (N/C) mice were fed Dox-containing food and painted with a 4-OH-Tam (Tam) solution (applied on the lower back skin; indicated with red rectangle) over a period of 10 days. For CPI treatment, experimental mice received four doses of αPD-1 and αCTLA-4 antibodies (“combo”). C) Skin pathology developed in the area of Ag expression in mice receiving Dox/Tam and combo CPIs. Distribution of clinical severity scores by treatment group is shown in pie charts, median is shown in dot plot. ** *P* = 0.0023 and ns = not significant by *t* test. n = 6 or 7, representative of 3 independent experimental repeats. D) Top: H&E (left) and CD3e staining by IHC of skin sections (20x) from representative mice in the indicated experimental group. Arrows highlight CD3e-positive cells. E) Quantification of epidermal thickness from mice in D, average ± standard deviation is shown. **** *P* < 0.0001 and ns = not significant by *t* test. n = 3 data points/mouse, representative of 3 independent experimental repeats. F) Representative H&E staining of skin sections from biopsies of healthy subjects (top) or patients with cutaneous lichenoid irAEs (bottom) (10x). G) Representative pictures of patients with cutaneous lichenoid irAEs (top) and comparison of macroscopic appearance of skin from a representative N/C mouse treated with Dox/Tam and combo CPIs (left) and a representative patient with a cutaneous lichenoid irAE (right). H) Normalized total counts of CD8 T cells (top) and CD4 T cells (bottom) purified from the skin of mice treated as indicated. Average ± standard deviation is shown. * *P* = 0.0495 (No Ag no CPIs vs + Ag no CPIs) and 0.0437 (No Ag no CPIs vs + Ag + combo), and ns = not significant by *t* test. n = 4 or 5, representative of 3 independent experimental repeats. I) Representative pictures (top) and H&E staining (bottom) from mice treated with Dox/Tam for Ag induction in the area indicated by the red rectangle coupled with combo CPIs with or without CD4 or CD8 T cell-depleting antibodies or both. J) Top: epidermal thickness from mice in I. Dot plot shows average ± standard deviation with dotted line representing average epidermal thickness in the absence of Ag induction. **** *P* < 0.0001 and ns = not significant by *t* test. n = 3 data points/mouse, representative of 2 independent experimental repeats. Bottom: clinical severity score of mice in I. Dot plot shows median of scores. * *P* = 0.0123 and ns = not significant by *t* test. n = 3, representative of 2 independent experimental repeats.

**Figure 2.**
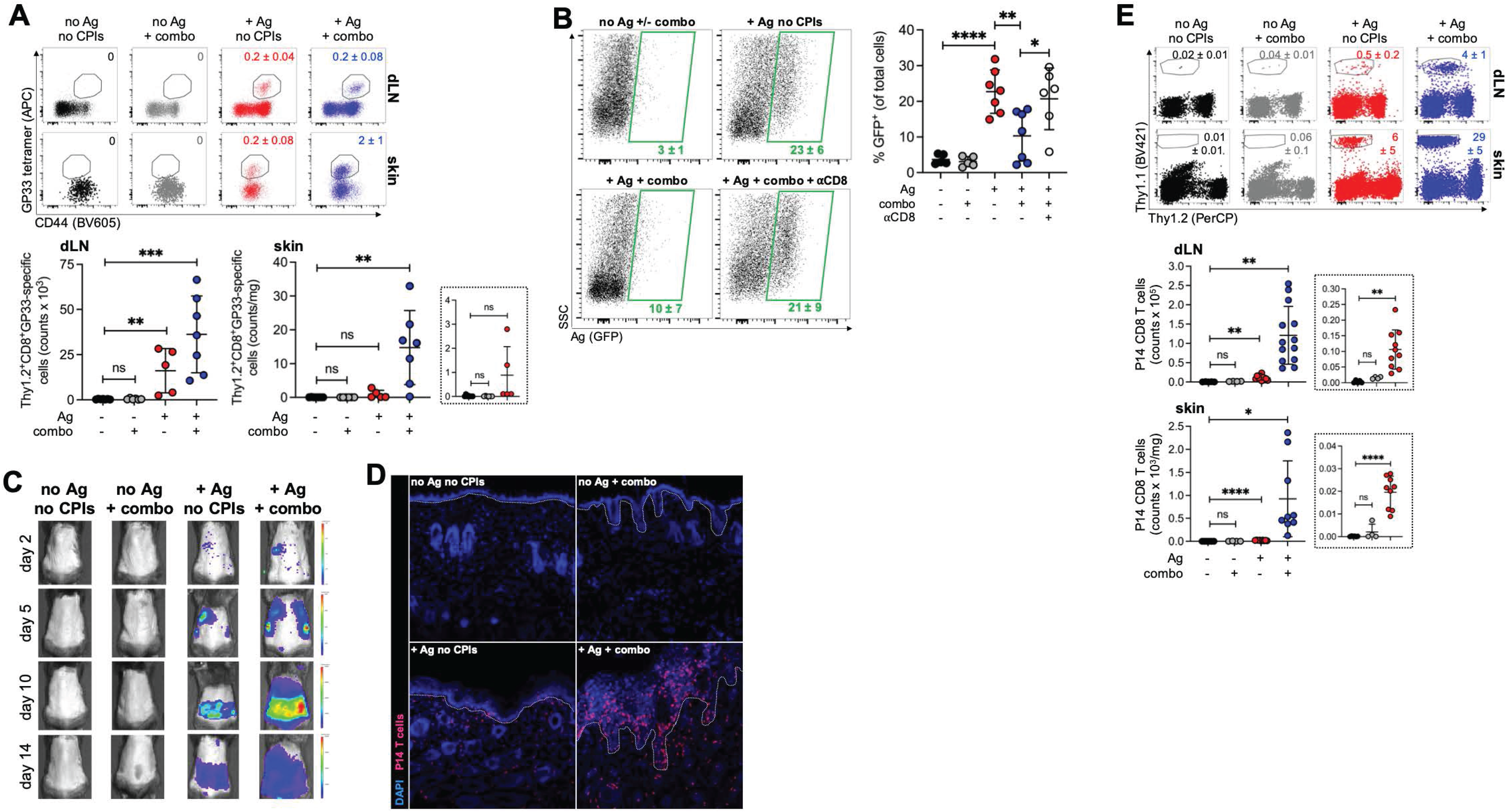
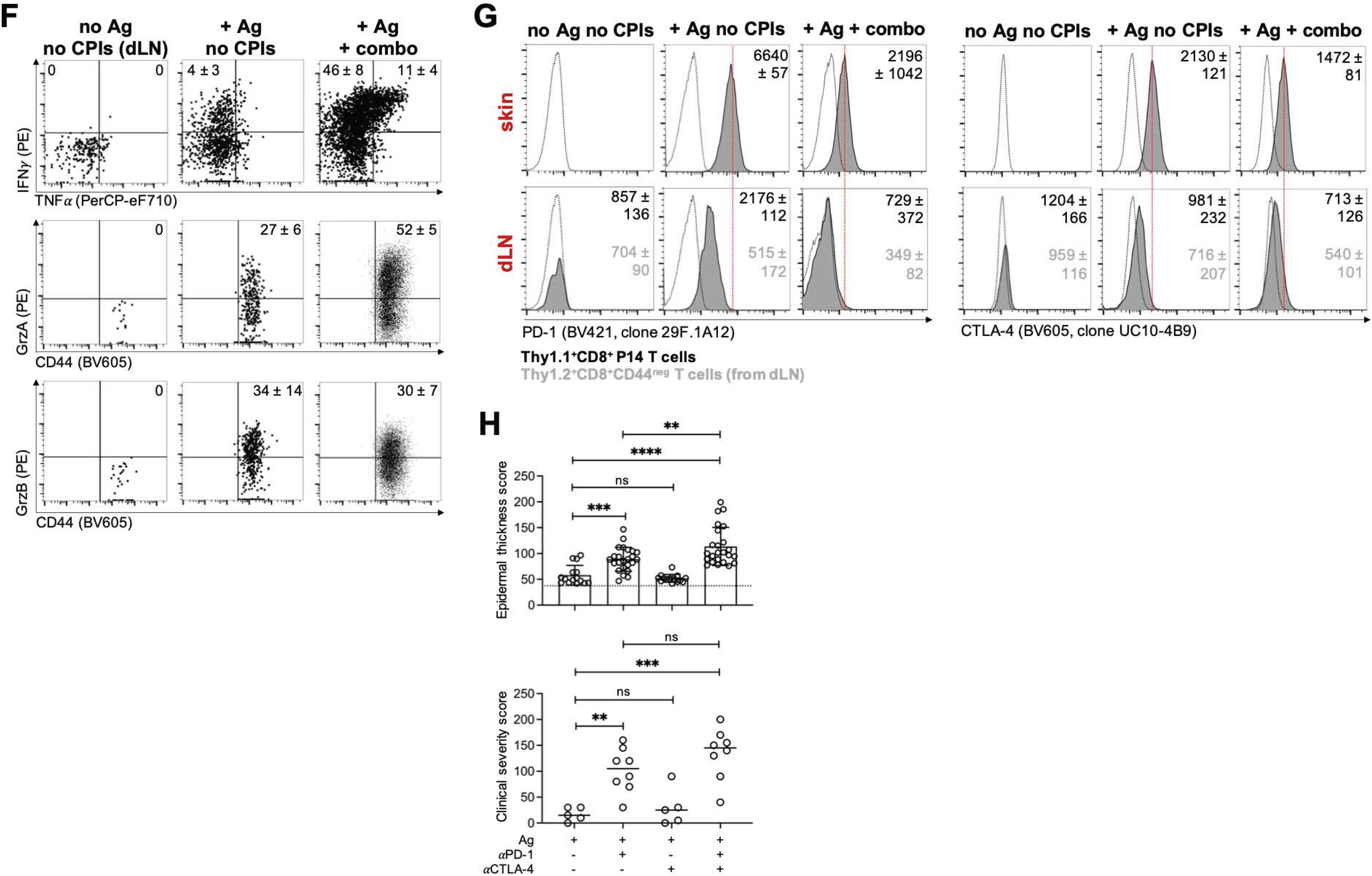
Ag-specific CD8 T cells acquire features of effector T cells but become pathogenic upon administration of CPIs. A) Frequency (top) and total counts (bottom) of endogenous GP33-specific CD8 T cells in local dLNs and Tam-treated skin from the indicated experimental groups. Numbers in facs plots represent average ± standard deviation of frequencies; average ± standard deviation is shown in dot plots. For dLNs, ** *P* = 0.006, *** *P* = 0.0008 and ns = not significant by *t* test; for skin, ** *P* = 0.004 and ns = not significant by *t* test. n = 5-7, representative of 3 independent experimental repeats. B) Representative flow cytometric analysis (left) and dot plot (right) of total skin cells harvested from mice treated as indicated and analyzed for GFP (Ag) expression. Numbers in facs plots indicate average ± standard deviation of frequencies of GFP+ skin cells. * *P* = 0.036, ** *P* = 0.0043 and **** *P* < 0.0001 by *t* test. n = 5-7, representative of 4 independent experimental repeats. C) IVIS imaging of fLuc-expressing P14 CD8 T cells in representative mice from the indicated experimental conditions and imaged after 2, 5, 10 and 14 days from the beginning of Dox/Tam dosing. Blue to red indicates increased signal intensity. n = 3, representative of > 3 experimental repeats. D) Confocal imaging of representative skin samples harvested at experimental end point from Tam-treated areas of mice from the indicated experimental conditions. Dotted line represents epidermis-dermis interface. Blue = DAPI, Magenta = dsRED (P14 CD8 T cells). n = 3. E) Frequency (top) and total counts (bottom) of transgenic Thy1.1/1.1+ P14 CD8 T cells in skin dLNs and Tam-treated skin from the indicated experimental groups. Numbers in facs plots represent average ± standard deviation of frequencies; average ± standard deviation is shown in dot plots. For dLNs, ** *P* = 0.0032 (No Ag no CPIs vs Ag no CPIs) and 0.0028 (No Ag no CPIs vs Ag + combo), and ns = not significant by *t* test; for skin, * *P* = 0.018, **** *P* < 0.0001, and ns = not significant by *t* test. n = 4-12, representative of 3 independent experimental repeats. F) Flow cytometric analysis of Thy1.1/1.1+ P14 CD8 T cells harvested from the Tam-treated skin (or skin dLNs of No Ag controls) of mice from the indicated experimental conditions. Cells were *ex vivo* stimulated with GP33-41 peptide prior to IFNγ and TNFα staining, peptide stimulation was not performed prior to GrzA/B staining. Numbers in facs plots indicate average ± standard deviation of frequencies of gated populations. n = 5, representative of 3 experimental repeats. G) Flow cytometric analysis of Thy1.1/1.1+ P14 CD8 T cells harvested from the skin and local dLNs of mice treated as indicated. Naïve Thy1.2^+^CD8^+^CD44^neg^ cells from skin dLNs were used as negative controls. Numbers in histograms represent average ± standard deviation of MFI. n = 5, representative of 3 experimental repeats. H) Epidermal thickness (top) and clinical severity score (bottom) from the skin of mice treated as indicated. Top: average ± standard deviation with dotted line representing average epidermal thickness in the absence of Ag induction. ** *P* = 0.0077, *** *P* = 0.0001, **** < *P* = 0.0001, and ns = not significant by *t* test. n = 3 data points/mouse, representative of 2 experimental repeat. Bottom: median of clinical severity scores is shown. ** *P* = 0.0013, *** *P* = 0.0003, and ns = not significant by *t* test. n = 5-8, representative of 2 experimental repeat.

### CPI therapy coupled with skin-specific Ag expression leads to CD8 T cell-mediated localized tissue pathology with features of human cutaneous lichenoid irAEs

To determine if the lack of pathology in N/C mice was due to the function of ICRs, we decided to concomitantly treat N/C mice with Dox/Tam and a combination of PD-1- and CTLA-4-specific blocking antibodies (Ag + combo) (**Fig 1B**). Between day 10-15, Ag + combo mice developed macroscopic tissue pathology requiring euthanasia of the animals by day 15 (**Fig 1C**). Notably, tissue pathology in these mice was highly restricted to the area of the skin where Ag expression was induced (indicated by the red rectangle, **Fig 1C**). To quantify the macroscopic pathology, we established a cutaneous clinical severity scoring system (see Methods section for details), and mice were scored blindly by a trained dermatologist. Skin pathology was characterized by varying degrees of erythema, lichenification and erosion/ulceration, with a median clinical severity score of 90 (**Fig 1C** and **S1**). By contrast, no significant increase in clinical severity score was observed in negative control mice (B6 or NINJA treated with Dox/Tam or untreated N/C mice) with or without CPIs (No Ag or No Ag + combo mice, **Fig 1C**). Histological analysis of the Tam-treated skin revealed significant changes in the epidermis and dermis of Ag + combo mice compared to all other experimental conditions (**Fig 1D**). Specifically, there was a significant increase in epidermal thickness (**Fig 1E**), and this was accompanied by a robust superficial inflammatory infiltrate primarily composed of lymphocytes, especially CD3e+ T cells (**Fig 1D**). Moreover, there was a visible concentration of T cells at the dermal-epidermal junction and within the epidermis that was accompanied by focal spongiosis and individual necrotic keratinocytes (**Fig 1D**). By contrast, no changes were observed in No Ag + combo mice (CPI alone), while T cells could be found scattered throughout the dermal layer of Ag-expressing mice without CPIs.

The macroscopic appearance and histology of Ag + combo N/C mice were reminiscent of cutaneous lichenoid irAEs that are found in patients after CPI treatment. Cutaneous irAEs are the most common irAEs, and lichenoid irAEs account for ~20% of these events (Schaberg et al.; Shi *et al.*). Histopathology of these cutaneous lichenoid irAEs typically demonstrates a superficial lichenoid infiltrate primarily composed of lymphocytes, with necrosis of epidermal keratinocytes and variable hyperkeratosis. To make a more direct comparison, we obtained biopsies from four patients who were clinically and pathologically diagnosed with lichenoid irAEs following CPI immunotherapy (**Fig 1F, G**). The clinical and histological features of human lichenoid irAEs were consistent with our clinical and histological findings in Ag + combo mice (**Fig 1D, E**). Therefore, in the presence of Ag expression in the skin, combined CPI treatment in N/C mice led to significant structural and pathological tissue changes and visible T cell infiltration of the epidermis with features of human cutaneous lichenoid irAEs.

We next asked whether tissue-infiltrating T cells had a pathogenic role in the development of localized lichenoid skin pathology in our model. To address this question, we first quantified total endogenous CD4 and CD8 T cells in the skin of experimental mice. We found that both populations were increased in Ag-expressing mice with and without CPIs, but only CD8 T cell counts were significantly different from No Ag (without CPI) controls, with CD8 T cells/mg skin quantified at 26 ± 19 (4-fold increase) in Ag mice and 61 ± 47 (10-fold increase) in Ag + combo mice (**Fig 1H**). Next, we depleted CD4 and/or CD8 T cells to determine if they were necessary for disease. We observed that depletion of CD8 T cells, but not of CD4 T cells, prevented the development of macroscopic skin pathology and histological changes in Ag + combo N/C mice (**Fig 1I, J**). Together, these data demonstrated that N/C mice could be used as a faithful model of human cutaneous lichenoid irAEs, in which disease was dependent on skin-specific Ag induction, CPI treatment, and CD8 T cells. Moreover, this provided a unique tool for determining how CPIs altered the biology of tissue-infiltrating skin-specific CD8 T cells and the mechanisms by which ICRs prevent disease.

### CPIs increase the effector functions of skin-infiltrating Ag-specific CD8 T cells

To better understand the impact of CPIs on skin-specific CD8 T cells, we analyzed endogenous GP33-specific CD8 T cells in Tam-treated skin and local dLNs. Ag induction alone led to an increase in the frequency of GP33-specific CD8 T cells (identified using GP33-loaded MHC class I tetramers) in the dLNs and Ag-expressing skin, from 0 to 0.2 ± 0.04% and 0.2 ± 0.08% of total CD8 T cells, respectively (**Fig 2A**). Moreover, Ag + combo treatment led to a significant increase of GP33-specific CD8 T cells compared to Ag alone, both in dLNs (36 ± 21 x 10^3^; 2-fold increase) and Ag-expressing skin (15 ± 10/mg; 15-fold increase) (**Fig 2A**).

Next, we determined if CPI administration could affect expression of self Ag in the skin. As GP_33-43_ is contained within the GFP protein, we used flow cytometric analysis of GFP to determine which cell types were expressing Ag at the experimental endpoint. GFP expression was detected in 23 ± 6% of total skin cells following Dox/Tam treatment of N/C mice (**Fig 2B**). More than 80% of GFP-expressing cells in Dox/Tam-treated N/C mouse skin were identified as EpCAM+CD45-, indicating that most of the cells expressing Ag after Dox/Tam treatment in our model were epidermal cells, potentially keratinocytes (**Fig S2A-B**). However, in skin from Ag + combo mice, the frequency of total Ag-expressing cells fell to 10 ± 7%, with the most significant loss occurring amongst the EpCAM+CD45-GFP+ cells (**Fig 2B**). This frequency was similar to baseline levels of GFP signal (autofluorescence) detected in negative controls (27 ± 10% of EpCAM+CD45− cells, **Fig S2A-B**). The expression of self Ag primarily by epidermal cells in the skin and the loss of these cells after CPI treatment is consistent with the increased infiltration of CD8 T cells into the epidermis after CPI treatment (**Fig 1H**) and the hypothesis that CPIs are driving loss of CD8 T cell tolerance towards Ag-expressing skin cells. In line with this, CD8 T cell depletion in Ag + combo-treated N/C mice rescued GFP expression in the skin (**Fig 2B** and **Fig S2A-B**), demonstrating that CD8 T cells were necessary to eliminate self Ag-expressing epidermal cells after combo CPI therapy.

To analyze the kinetics, location, phenotypes, and functions of GP33-specific CD8 T cells in our model, we bred Thy1.1/1.1+ GP33-specific TCR transgenic P14 mice (Pircher et al.) to mice expressing firefly Luciferase (fLuc) or mice expressing the fluorescent protein dsRED, which allowed us to adoptively transfer 5 x 10^3^ naïve fLuc+ or dsRED+ P14 CD8 T cells into our mouse models to track Ag-specific CD8 T cells responding to Ag or Ag + combo treatment in N/C mice (**Fig S3A**). The transfer of naïve P14 CD8 T cells led to more severe localized skin pathology in Ag + combo mice than in mice without P14 CD8 T cell transfer (increase in median clinical severity score from 90 to 140), and also led to an increase in the median clinical severity score of Ag alone mice (from 2.5 to 25), which was significantly different from No Ag controls, but far below the Ag + combo CPI-treated mice without adoptive transfer of P14 CD8 T cells (**Fig S3B-C**). This is in line with the fact that GP33-specific P14 CD8 T cells express higher-avidity Ag-specific TCRs and were present at higher precursor frequencies than endogenous GP33-specific CD8 T cells. Because of this, where possible, we confirmed that our subsequent findings regarding the functions and phenotypes of P14 CD8 T cells were also observed in endogenous GP33-specific CD8 T cells.

To track the kinetics of the GP33-specific CD8 T cell response *in vivo* in N/C mice, we used adoptive transfer of naïve fLuc-expressing P14 CD8 T cells, and imaged animals by IVIS starting at day 2 after initiating Dox/Tam treatment. Initial expansion of skin Ag-specific CD8 T cells was observed in Tam-treated skin dLNs and spleens between day 2 and 5 in Dox/Tam-treated N/C mice, and by day 10, fLuc+ P14 CD8 T cells had accumulated in the Ag-expressing skin area, but not beyond (**Fig 2C**). The early infiltration of P14 CD8 T cells was consistent with the onset of macroscopic changes in skin observed in Ag + combo mice (**Fig 1C**), but early and sustained infiltration of P14 CD8 T cells also occurred in mice with Ag only, suggesting that disease after CPI treatment was not simply a result of T cell infiltration into skin. To assess the location of GP33-specific CD8 T cells within the skin after Ag induction, we transferred naïve dsRED-expressing P14 CD8 T cells and analyzed skin from mice 15 days after initiation of Dox/Tam. Confocal microscopy confirmed that dsRED+ P14 CD8 T cells did not infiltrate the skin in No Ag control mice, despite Dox/Tam or CPI treatment (**Fig 2D**). In mice where Ag was expressed in the skin, P14 CD8 T cells were present, but these cells were mostly localized to the dermis (**Fig 2D**), consistent with the scattered distribution of CD3e+ T cells observed in Ag-expressing N/C mice by IHC staining (**Fig 1D**). By contrast, robust infiltration of P14 CD8 T cells into the epidermis was observed in Ag + combo mice (**Fig 2D**), which was consistent with the IHC analysis of CD3e+ T cells in mice without P14 CD8 T cells (**Fig 1D**). Thus, skin Ag-specific CD8 T cells infiltrated into the sites where self Ag was expressed after induction, and their ability to infiltrate the epidermis was enhanced by CPI treatment. The latter is in line with the fact that GFP+ epidermal cells were eliminated in Ag + combo mice (**Fig S2**).

Flow cytometric analysis confirmed that expansion and accumulation of Thy1.1/1.1+ P14 CD8 T cells was detected in Tam-treated skin and local dLNs following Ag induction (0.5 ± 0.2% and 6 α 5% of total cells in dLNs and skin, respectively), and this was significantly increased in Ag + combo mice (4 ± 1% and 29 ± 5% of total cells in dLNs and skin, respectively, **Fig 2E**). To assess the function of skin-infiltrating P14 CD8 T cells, we analyzed their expression of cytolytic granule proteins (Granzymes A and B; GrzA/B) and their ability to produce pro-inflammatory cytokines after *ex vivo* peptide stimulation. As negative controls, we analyzed P14 CD8 T cells from the skin dLNs of Dox/Tam-treated B6 or NINJA control mice (No Ag controls). The P14 CD8 T cells in No Ag mice were GrzA- and GrzB-negative and did not produce IFNγ or TNFα after *ex vivo* stimulation with GP33-41 peptide (**Fig 2F**). This is in line with P14 CD8 T cells remaining naïve in No Ag mice. By contrast, many P14 CD8 T cells in Ag mice produced GrzA (27 ± 6%) and GrzB (34 ± 14%), and 4 ± 3% of these P14 CD8 T cells had the capacity to produce IFNγ (but not TNFα). The functions of skin P14 CD8 T cells in Ag + combo mice were even further increased, as a higher fraction of P14 CD8 T cells produced GrzA (52 ± 5%) and IFNγ (46 ± 8%), with a notable increase in IFNγ+ TNFα+ double producers (11± 4%). We did not observe increased frequency of GrzB+ P14 CD8 T cells in Ag + combo mice (30 ± 7%, **Fig 2F**), which was unexpected, as GrzB is a key cytolytic effector protein in CD8 T cells. These data highlight how CPI treatment enhanced the production of several effector proteins by skin Ag-specific CD8 T cells. Moreover, they reveal that many skin-infiltrating Ag-specific CD8 T cells also acquired effector functions in mice that expressed Ag in skin without treatment with CPIs. These phenotypes differed from T cells in conventional models of anergy or deletional tolerance, where cells do not acquire effector functions and typically fail to infiltrate tissues. Similar effector functions were also observed in endogenous GP33-specific CD8 T cells in skin (**Fig S4**).

### PD-1 blockade but not CTLA-4 blockade is sufficient to drive cutaneous lichenoid irAEs in N/C mice

To better understand how CPI treatment impacted GP33-specific CD8 T cells we analyzed the expression of PD-1 and CTLA-4 on Ag-specific CD8 T cells in Tam-treated skin and local dLNs. Compared to Ag-specific P14 CD8 T cells harvested from the dLNs, skin-infiltrating P14 CD8 T cells expressed higher levels of both PD-1 (MFI 6640 ± 57 vs 2176 ± 112) and CTLA-4 (MFI 2130 ± 121 vs 981 ± 232) following skin-specific Ag induction (**Fig 2G**). These expression levels were decreased in both sites after combined CPI treatment (**Fig 2G**). This decrease was likely the result of the *in vivo* administered CPI antibodies blocking the *ex vivo* staining of P14 CD8 T cells with anti-PD-1 and anti-CTLA-4 antibodies for flow cytometry, as the gene expression levels of *Pdcd1* and *Ctla4* were similar between P14 CD8 T cells with and without CPI treatment (**Fig S5**). This demonstrated that the blocking-antibodies targeting ICRs were bound to self-reactive CD8 T cells in the skin. We also noted that skin-infiltrating P14 CD8 T cells expressed several different ICRs transcriptionally, including *Lag3*, *Havcr2* (Tim-3), and *Tigit,* following localized Ag induction (**Fig S5**).

Cutaneous lichenoid irAEs in patients are more commonly observed in individuals treated with PD-1-blocking agents (Coleman *et al.*; Shi *et al.*), so we next determined if this was also the case in the N/C model. Skin-specific induction of Ag with blockade of PD-1 alone resulted in localized skin pathology similar to combined PD-1/CTLA-4 blockade, although with overall reduced severity (**Fig 2H**; epidermal thickness score 89 ± 23 vs 114 ± 37 and median clinical severity score of 105 vs 145). By contrast, there was no significant disease observed when Ag-expressing N/C mice were treated with anti-CTLA-4 alone (**Fig 2H**). These data demonstrate that disruption of PD-1 ICR function, potentially within the Ag-expressing skin, led to pathology by Ag-specific effector CD8 T cells. This is consistent with the notion that ICRs play a critical role in preventing self Ag-specific CD8 T cell pathology, preserving peripheral tissue integrity and tolerance.

### Single-cell transcriptomic analysis of skin cells from No Ag, Ag, and Ag + combo mice

To understand the transcriptomic changes that occurred in the skin between Ag and Ag + combo N/C mice, and how these resulted in localized lichenoid tissue pathology, we performed single-cell RNA sequencing (scRNAseq) analysis of both hematopoietic and non-hematopoietic local skin cells from No Ag control mice, skin Ag-expressing mice and skin Ag-expressing mice treated with combined CPIs (**Fig 3A**). For this purpose, we adoptively transferred naïve dsRED+ P14 CD8 T cells into NINJA or N/C mice and treated them with Dox/Tam (for No Ag and Ag conditions, respectively), with a group of Dox/Tam-treated N/C mice also receiving combined anti-PD-1/CTLA-4 antibodies (Ag + combo; n = 2 mice for all groups). Note, No Ag mice also received dsRED+ P14 CD8 T cells and Dox/Tam, the same as the other experimental groups (to control for Dox/Tam-dependent effects that were not dependent on skin-specific Ag induction). We validated that, upon treatment, each experimental mouse used for this analysis showed the expected phenotype based on localized macroscopic skin pathology as well as functionality of Ag-specific CD8 T cells (**Fig S6**). Next, we processed the Tam-treated skin to isolate single cell suspensions that we subjected to scRNAseq with paired V(D)J sequencing. For data analysis, six total samples were co-embedded (2 No Ag, 2 Ag, and 2 Ag + combo) to perform Seurat-based clustering analysis and results were visualized using Uniform Manifold Approximation and Projection (UMAP) plots for dimension reduction (**Fig 3B**). This initial analysis showed four major cell populations across samples comprised of epithelial cells, fibroblasts, myeloid cells and T cells, and important differences in the fraction of cells in each major cell population were observed depending on experimental condition. For example, epithelial cells were mostly derived from No Ag and Ag samples, with relatively lower proportion of epithelial cells from Ag + combo samples (**Fig 3B-C**). This was in line with the significant loss of GFP+ EpCAM+CD45− cells observed by flow cytometry in Ag + combo N/C mice (**Fig 2B** and **S2**). In parallel, there was an increase in the representation of T cells and myeloid cells in samples from Ag and Ag + combo conditions (**Fig 3B-C**).

**Figure 3.**
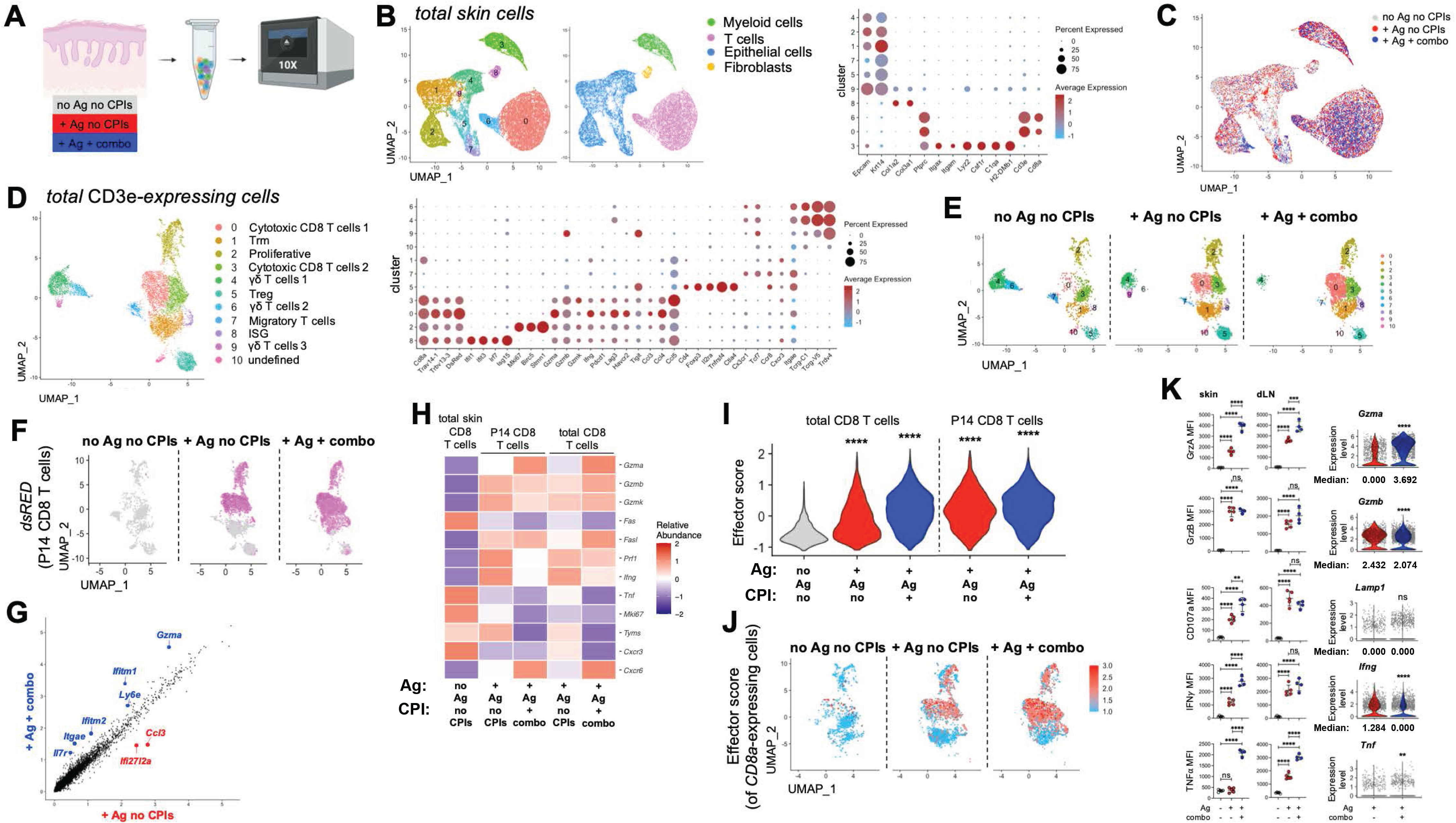
Transcriptomic analysis confirms transition to effector state by Ag-specific CD8 T cells following skin-specific Ag expression. A) Experimental workflow: total skin was harvested from the Tam-treated area of NINJA mice (“No Ag no CPIs”) and N/C mice in the absence or presence of combo CPIs (“+ Ag no CPIs” and “+ Ag + combo”, respectively). Single cell suspensions were processed for scRNAseq with paired V(D)J sequencing using the 10X Genomics platform. n = 10162 cells (total skin cells from “No Ag no CPIs” group), 9017 cells (total skin cells from “+ Ag no CPIs” group), 6019 cells (total skin cells from “+ Ag + combo” group). B) Identification of total skin cell clusters across all samples (left) and bubble plot of canonical marker genes utilized to identify each cluster (right). C) Distribution of skin cells across clusters colored based on experimental group. D) Sub-clustering of CD3e+ T cells from the dataset in B (left) and bubble plot of canonical marker genes utilized to identify each sub-cluster (right). E) Distribution of T cells across the clusters identified in D shown by experimental condition. F) Distribution of *dsRED*-expressing cells (P14 CD8 T cells) among total CD8 T cells sequenced is shown by experimental condition. n = 1203 (P14 CD8 T cells from “+ Ag no CPIs” group) and 2053 (P14 CD8 T cells from “+ Ag + combo” group). G) Scatterplot of average gene expression in P14 CD8 T cells from “+ Ag no CPIs” (x) and “+ Ag + combo” (y) experimental conditions. H) Gene expression heatmap of the genes included in the CD8 T cell effector score displayed by experimental condition. I) Distribution of CD8 T cell effector scores from the genes in H calculated for total CD8 T cells and P14 CD8 T cells from the indicated experimental conditions. **** *P* < 0.0001 by Wilcoxon rank sum test for comparisons of no “Ag no CPIs” vs “+ Ag no CPIs” or “+ Ag no CPIs” vs “+ Ag + combo”. n = 884 cells (total skin CD8 T cells from “No Ag no CPIs” group), 2201 cells (total skin CD8 T cells from “+ Ag no CPIs” group), 2565 cells (total skin CD8 T cells from “+ Ag + combo” group), 1203 (P14 CD8 T cells from “+ Ag no CPIs” group) and 2053 (P14 CD8 T cells from “+ Ag + combo” group). J) UMAP of CD8 T cell effector score computed for clustered CD8 T cells and displayed by experimental condition. K) Flow cytometric analysis and scRNAseq analysis of Thy1.1/1.1+ P14 CD8 T cells harvested from the skin and skin dLNs of mice treated as indicated (flow cytometry was performed after *ex vivo* stimulation with GP33-41 peptide). For MFI, dot plots show average ± standard deviation with *** *P* = 0.0009 and **** *P* < 0.0001 (GrzA); **** *P* < 0.0001 and ns = not significant (GrzB); ** *P* = 0.0044, **** *P* < 0.0001 and ns = not significant (CD107a); **** *P* < 0.0001 and ns = not significant (IFNγ); **** *P* < 0.0001 and ns = not significant (TNFα) by *t* test. n = 4-5, representative of 3 experimental repeats. For violin plots of gene expression level distributions ** *P* = 0.01 and **** *P* < 0.0001 by Wilcoxon rank sum test. n = 1203 (P14 T cells from “+ Ag no CPIs” group) and 2053 (P14 T cells from “+ Ag + combo” group).

### Minimal Transcriptional differences in Ag-specific CD8 T cells in skin with and without checkpoint immunotherapy

To gain a better understanding of the transcriptional differences between T cells across the experimental conditions analyzed, we first sub-clustered all CD3e+ cells sequenced in our experimental samples (**Fig 3D-E**). This analysis revealed 10 T cell clusters, with substantial differences observed, particularly between the No Ag and Ag experimental groups (**Fig 3E**). Expression of Ag in the skin was associated with increased abundance of T cells in clusters 0, 1, and 2, which were characterized by cytotoxic CD8 T cell, tissue-resident memory T cell (Trm) and proliferative transcriptional profiles, respectively (**Fig 3D-E**). As Ag-specific CD8 T cells in this experiment expressed dsRED, we confirmed that *Dsred* transcripts were largely found in T cells from clusters 0-3 in Ag-expressing mice (**Fig 3D, F** and **S11A**). These Ag-driven increases were paralleled by decreased representation of cells from clusters 4, 6, and 9, which were primarily comprised of cells with a γδ T cell gene expression profile (**Fig 3D-E**). Here, it is worth noting again that No Ag and Ag mice were both treated with Dox/Tam and only differed in their capacity to express Ag in skin cells after treatment. These data are consistent with the idea that local infiltration of CD8 T cells following skin-specific Ag expression led to alterations in the local composition of T cell populations.

While many differences were identified comparing T cells in No Ag vs Ag samples, surprisingly few differences were found when comparing the transcriptional profiles of T cells from Ag and Ag + combo samples. Here, the most notable differences between Ag vs. Ag + combo were a further decrease in the γδ T cell clusters and cluster 7 (Treg gene profile), and an increased abundance of cytotoxic CD8 T cell clusters (**Fig 3D-E**).

We next focused on the transcriptional differences between *Dsred*-expressing P14 CD8 T cells from the Ag and Ag + combo experimental conditions. A scatterplot showing average gene expression levels of P14 CD8 T cells from these two conditions unexpectedly revealed only a handful of differentially expressed genes that included increases in *Gzma*, *Ifitm1* and *Ifitm2*, *Ly6e*, *Itgae* and *Il7r,* and decreases in *Ccl3* and *Ifi27l2a* in Ag + combo mice (**Fig 3G**). Similar results were seen when comparing Ag and Ag + combo P14 CD8 T cells within clusters 0-3 (**Fig S11B**). This was surprising, as we expected significant differences in effector gene expression between Ag-specific CD8 T cells in Ag and Ag + combo conditions based on the observed pathology and functional differences in these mice (**Fig S6**). To better quantify this, we focused on a panel of genes encoding CD8 T cell effector molecules, including *Gzma*, *Gzmb*, *Gzmk*, *Prf1* and *Ifng* (**Fig 3H**) and created a CD8 T cell “effector score”. This allowed for a direct comparison of skin-infiltrating CD8 T cells from the different samples. Overall, there was a sharp increase in effector score between CD8 T cells (total and Ag-specific) from the No Ag to Ag conditions, but the effector score increase was less apparent, although statistically significant, between Ag and Ag + combo samples (**Fig 3I**). Moreover, this increase was driven by changes in *Gzma*, as a calculation of effector score without this gene showed significantly <u>higher</u> effector scores in P14 T cells in Ag vs. Ag + combo (**Fig S11C**). A pseudo-colored UMAP plot of effector score on clustered CD8 T cells showed that the cells with the highest effector scores were located in clusters 0-3, and that these cells were present in both Ag and Ag + combo conditions (**Fig 3J**). However, despite similar (or higher) gene expression levels for most effector genes (**Fig 3H**), flow cytometric analysis confirmed significant increases in the skin-infiltrating P14 CD8 T cells for protein expression of GrzA, GrzB, surface CD107a (marker of degranulation capacity), IFNγ and TNFα in Ag + combo mice (**Fig 3K**; Note, we have quantified median protein (left) and mRNA (right) expression). When comparing skin-infiltrating and skin dLN-derived P14 CD8 T cells from Ag only mice, we noticed that in the skin, cells had lower protein expression levels of IFNγ (MFI 1208 ± 220 vs 2100 α 394), reduced surface staining of CD107a (MFI 205 ± 33 vs 480 ± 82) and lacked TNFα (**Fig 3K**). By contrast, GrzB, surface CD107a, and IFNγ were similar in dLN CD8 T cells in Ag vs Ag + combo (**Fig 3K**). These data were consistent with a model where skin-specific Ag expression in N/C mice was sufficient to drive the development of Ag-specific CD8 T cells with high effector capacity (at the transcriptional level), but that their effector functions were post-transcriptionally suppressed upon migration into the skin. Here, we postulate that CPIs allowed self Ag-specific CD8 T cells to produce effector proteins and mediate pathogenesis.

### Low-penetrance spontaneous skin pathology in Ag-expressing non-CPI treated N/C mice correlates with highly functional Ag-specific CD8 T cells

Treatment of N/C mice with Dox/Tam and combined CPIs resulted in lichenoid skin pathology with near 100% penetrance and disease was more severe when P14 CD8 T cells were present (**Fig 1C** and **S3**). Moreover, under these experimental conditions there was a 100% correlation between the development of skin pathology and the presence of highly functional IFNγ+TNFα+ Ag-specific CD8 T cells (**Fig 2F**). However, through careful analysis of dox/tam-treated Ag mice (without transferred P14 CD8 T cells), we noticed that ~17% of these mice developed mild skin pathology (clinical severity score = 16 - 50) and had IFNγ+TNFα+ Ag-specific CD8 T cells (**Fig S7**). The penetrance of disease was increased in Ag-expressing N/C mice with P14 CD8 T cells, but the 100% correlation with presence of IFNγ+TNFα+ Ag-specific CD8 T cells was maintained (**Fig S7**). These data are consistent with the idea that CPI treatment did not endow self Ag-specific CD8 T cells with unique pathogenic T cell effector functions, but rather it lowered the threshold for those cells to achieve that pathogenic state.

### Ag-expressing epidermal cells in skin do not protect themselves by upregulating PD-L1

Because skin-specific Ag induction coupled with systemic PD-1 blockade in our model was sufficient to drive CD8 T cell-dependent skin destruction, we investigated which skin cell types expressed PD-L1 following Ag expression in the skin. We hypothesized that epidermal cells themselves would express PD-L1, as EpCAM+CD45− skin cells were the Ag-expressing cell type and these cells were eliminated by CD8 T cells in the context of CPI therapy (**Fig S2**). However, neither analysis of *Cd274* (PD-L1) expression by epithelial cells nor flow cytometric analysis of EpCAM+CD45− skin cells showed that these cells expressed PD-L1 following localized Ag induction (**Fig S8A-B**). This was true even when we analyzed just the population of total skin Ag-expressing cells (**Fig S8C**). Of note, we also did not observe significantly increased expression of *Cd274*/PD-L1 by GFP negative EPCAM+ cells in Ag + combo mice by scRNAseq or flow cytometry, but the few remaining Ag-expressing (GFP+) EPCAM+ cells skin cells did show increased PD-L1 protein expression following combined CPI treatment (**Fig S8A-C**).

PD-L1 is a known target downstream of IFNγ receptor signaling (Garcia-Diaz et al.) and skin Ag-specific CD8 T cells in our model were capable of producing some IFNγ protein (**Fig 2F** and **3K**). We therefore analyzed the expression levels of multiple genes, including *CD274*, that have been shown to be upregulated as a consequence of IFNγ-dependent signaling (Ayers et al.; Benci et al.; Combes et al.; Liu et al.; Luoma *et al.*; Szabo et al.) (see Methods section for details) and developed an “Interferon-stimulated gene score” or ISG score. Few skin epithelial cells in Ag + combo mice were ISG score signature-positive compared with No Ag or Ag only mice (**Fig S8D**). This may be explained by the fact that skin epithelial cells in Ag only and Ag + combo mice had low *Ifngr1* and *Ifngr2* expression (**Fig S9**). Furthermore, these data suggest that PD-L1 expression on epidermal cells is not regulating the pathogenicity of Ag-specific CD8 T cells in skin from Ag only mice.

### Infiltration of Ag-specific CD8 T cells into Ag-expressing skin alters the landscape of local myeloid cells and leads to accumulation of PD-L1+ CD11b+CD11c+ cells

Through our scRNAseq analysis, we noticed that *Cd274* transcripts and ISG score were upregulated by myeloid cells following skin-specific Ag induction (**Fig S8A, D**). Sub-clustering of skin-derived myeloid cells revealed that *Cd274* was specifically upregulated by two clusters altogether characterized by high expression of *Itgax* (CD11c), *Itgam* (CD11b), *C1qa*, *H2DMb1*, *Lyz2* (LysM) and *Csf1r* (**Fig 4A-B** and **S10**). Moreover, while the representation of cells in clusters 1 (Langerhans cells) and 3 (CD103+ DCs) was greatly changed by local induction of Ag expression, most of these cells remained low for *Cd274* transcripts (**Fig 4A-B**). Analysis of total skin cells by flow cytometry revealed that CD45+CD11b+CD11c+ cells had significant increases in frequency (17.2 ± 5% vs. 7.5 ± 1% of CD45+ cells) and PD-L1 protein expression (MFI: 2594 ± 372 vs 1229 ± 531) in Ag-expressing compared to No Ag mice, respectively (**Fig 4C-D**). Of note, Ag expression was not detected in any of the CD45+ populations from Ag-expressing N/C mice (**Fig 4C**), in line with Ag induction occurring primarily in skin epidermal cells in our model.

**Figure 4.**
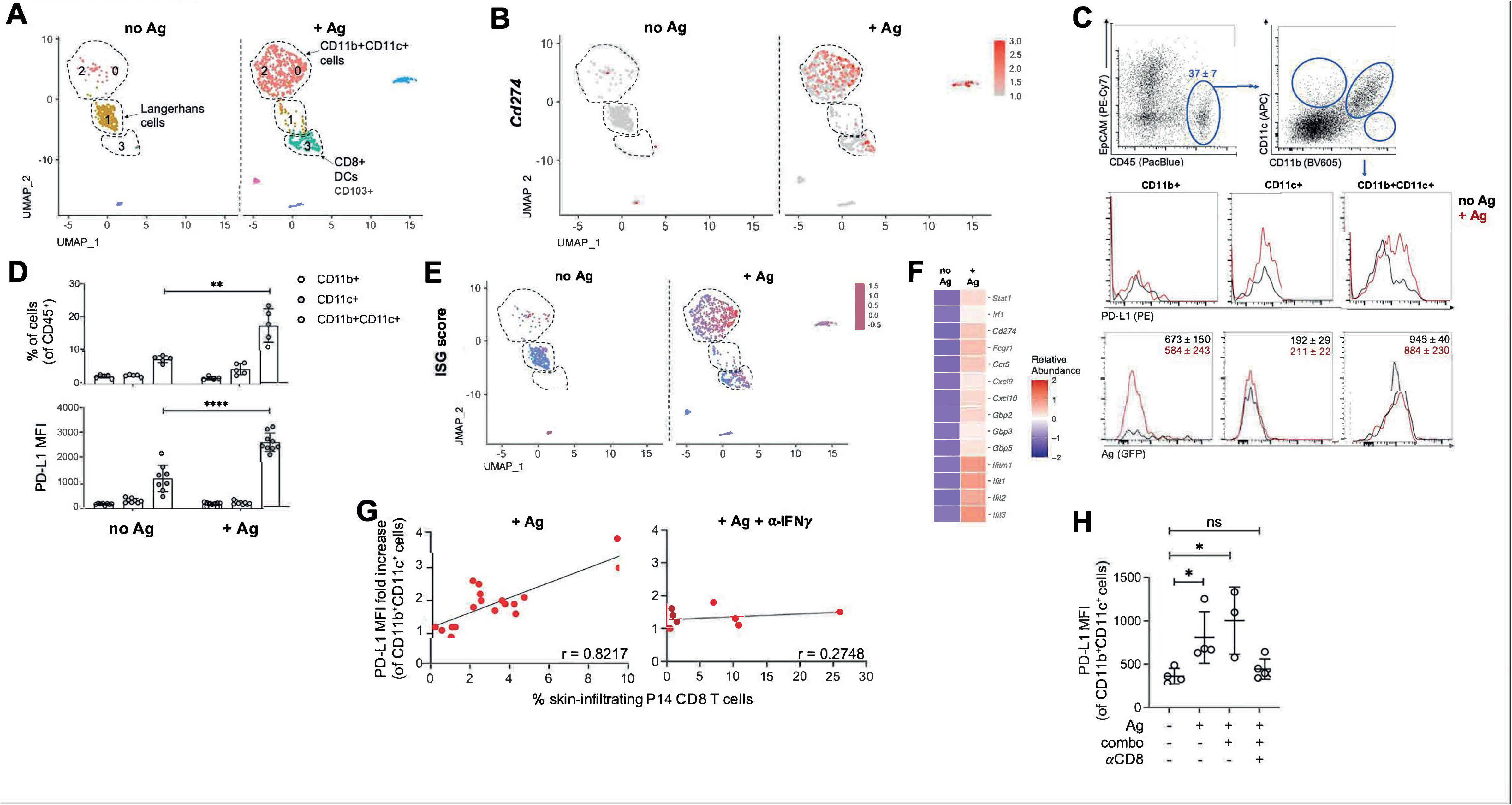
Skin CD11b+CD11c+ cells upregulate an IFN*γ*-dependent gene signature and PD-L1 expression in response to Ag induction and local infiltration of Ag-specific CD8 T cells. A) Sub-clustering of skin myeloid cells from the dataset in Fig 3B shown by experimental condition. B) Expression of *CD274* (PD-L1) across myeloid cells from A and shown by experimental condition. C) CD45+EpCAM− cells from skin are analyzed for CD11b and CD11c expression and the indicated populations are analyzed by PD-L1 and Ag (GFP) expression. Numbers in EpCAM/CD45 dot plot indicate average ± standard deviation of the frequency of CD45+EpCAM− cells across the No Ag/+ Ag experimental conditions. Numbers in GFP histograms show average ± standard deviation of GFP MFI. n = 5, representative of 3 experimental repeats. D) Frequency (top) and PD-L1 MFI (bottom) of the indicated CD45+ populations harvested from the skin of mice treated as shown and analyzed by flow cytometry. ** *P* = 0.0032 and **** *P* < 0.0001 by *t* test. n = 5-9, representative of 3 experimental repeats. E) Hallmark gene set score for IFN-stimulated genes (“ISG score”) computed for myeloid cells from A and displayed by experimental condition. F) Expression heatmap of genes included in the ISG score calculated for skin CD11b+CD11c+ cells from A and E and shown by experimental condition. G) Correlation analysis between frequency of skin-infiltrating P14 CD8 T cells (x) and PD-L1 MFI of CD11b+CD11c+ skin cells (y) from N/C mice treated with Dox/Tam (+ Ag) or with Dox/Tam and IFNγ-blocking antibody (+ Ag + α-IFNγ). For + Ag condition, slope is ≠ 0 with *P* < 0.0001; for + Ag + α-IFNγ condition, slope is not different from 0 with *P* = 0.4743 by simple linear regression analysis. n = 8-18, from 2 experimental repeats. For each experiment, PD-L1 MFI fold increase was calculated as a ratio between the PD-L1 MFI of CD45+CD11b+CD11c+ skin cells in the indicated experimental condition and the average PD-L1 MFI of CD45+CD11b+CD11c+ skin cells in “No Ag” controls. H) PD-L1 MFI of CD45+CD11b+CD11c+ cells harvested from the skin of mice treated as indicated. Dot plot shows average ± standard deviation with * *P* = 0.0284 (untreated vs + Ag) or 0.0215 (untreated vs + Ag + combo), and ns = not significant by *t* test. n = 5, representative of 3 experimental repeats.

Skin-specific Ag induction in N/C mice was paralleled by increased ISG score by CD11b+CD11c+ cells (**Fig 4E, F**). Our transcriptomic analysis confirmed that skin-infiltrating CD8 T cells were the only cells to express *Ifn* and no other transcripts for type I or type III *Ifn* family members were detected in any cells (not shown). This suggested that upregulation of PD-L1 and other IFNγ-dependent genes by skin CD11b+CD11c+ cells was a consequence of local infiltration of Ag-specific CD8 T cells following Ag induction. In fact, a strong correlation was found between frequency of skin-infiltrating P14 CD8 T cells and PD-L1 MFI of skin CD11b+CD11c+ cells in Ag-expressing N/C mice (r = 0.8217, **Fig 4G**). This correlation was lost when IFNγ-blocking antibodies were administered during skin-specific induction of Ag (r = 0.2748, **Fig 4G**). Moreover, we also observed that skin CD11b+CD11c+ cells failed to significantly upregulate protein expression of PD-L1 in CD8 T cell-depleted N/C mice (**Fig 4H**). These data indicate that, in our model, local release of IFNγ by skin-infiltrating effector-like Ag-specific CD8 T cells led to increased PD-L1 expression by skin CD11b+CD11c+ cells.

## Discussion

ICRs have been shown to promote peripheral T cell tolerance by driving deletion or anergy of self-reactive CD8 T cells following priming by tolerogenic DCs (Greenwald et al.; Keir *et al.*; Probst *et al.*; Probst et al.; Redmond *et al.*; Tivol *et al.*). This mechanism should lead to the absence of self-reactive CD8 T cells with pathogenic potential in healthy individuals, but the frequent occurrence of irAEs in cancer patients following CPI immunotherapy suggests that ICRs may have an active role in preventing immune-mediated pathology from self-reactive T cells. In a novel animal model, we tested the impact of *de novo* self Ag expression in skin and found that this did not lead to T cell anergy/deletion, but rather to infiltration of skin by self Ag-specific CD8 T cells with an effector-T-cell-like transcriptional program. Skin-infiltration by CD8 T cells drove compensatory changes in the local myeloid compartment via IFNγ, including significant increases PD-L1+ CD11b+CD11c+ cells. Yet, despite high effector potential and residence a self-antigen-expressing tissue for several days, the self-antigen-specific CD8 T cells did not cause tissue pathology. Tissue integrity was disrupted upon blockade of PD-1 or PD-1/CTLA-4, which led to CD8 T cell-mediated immunopathology directed against Ag-expressing epidermal cells. This recapitulated the cardinal features of cutaneous lichenoid irAEs in CPI-treated patients. Importantly, despite the dramatic change in CD8 T cell functions and pathological outcome with CPIs, there were relatively few transcriptional changes observed in tissue-infiltrating CD8 T cells, highlighting the importance of ICRs in keeping skin-specific CD8 T cells from becoming pathogenic in healthy skin. Together, these data show that ICRs can actively regulate self-reactive CD8 T cells and this mechanism maintains tolerance towards healthy skin. Disruption of this T cell-extrinsic regulation by CPIs unleashes the effector functions of self-reactive CD8 T cells, leading to pathologic tissue destruction and providing a potential mechanism for how CPI treatment leads to cutaneous lichenoid irAEs.

The pathogenic mechanisms of irAEs are poorly understood, but for many irAEs clinical evidence supports the hypothesis that self-reactive T cells specific for local tissue Ag cause localized tissue pathology following CPI administration (Johnson *et al.*; Luoma *et al.*; Shi *et al.*; Yasuda *et al.*). In this context, it is not yet clear when endogenous self-reactive T cells are exposed to their cognate Ag and where CPIs are acting to promote pathogenic activation of these cells. Two recently published studies have indicated that irAE pathogenesis correlates with presence of T cell populations within the target tissue. In one case, PD-1-blockade-dependent thyroiditis in mice was associated with thyroid Ag-specific central and effector memory T cells in both thyroid gland and thyroid dLNs prior to CPI administration (Yasuda *et al.*). In the other study, analysis of the lamina propria from patients with CPI-induced colitis showed expansion of effector-like CD8 T cells that were clonally related to cells with Trm-phenotype (Luoma *et al.*). Therefore, it is likely that Ag-experienced effector T cells can become resident memory cells in otherwise healthy tissues and yet remain in a non-pathogenic state. Here, the Trm could be specific for self-antigens or to other antigens that are presented in the tissue (*i.e.,* antigens from the microbiome). CPIs would then act to unleash the pathogenic capacity of these tissue-resident T cells. The data in our current study are consistent with this hypothesis and show the potential of skin-reactive T cells to mediate immunopathology after CPI treatment. Moreover, longer-term analyses of Ag only N/C mice suggest that Ag-specific CD8 T cells in skin can form Trm, and that the T cells in these mice also remain CPI-responsive (data not shown).

Most current irAE models rely on CPI treatment of autoimmune-prone or disease-induced mice (Perez-Ruiz et al.; Yasuda *et al.*), which does not allow for mechanistic understanding of how CPI treatment breaks physiological tolerance. The NINJA model used here uniquely allows one to study endogenous self-antigen-specific CD8 and CD4 T cell responses in an controlled manner, which allow one to determine how these T cells differ with and without CPI treatment. This will allow further investigations into mechanistic aspects of peripheral tolerance and will provide a platform for testing therapeutics to ameliorate irAEs or screen new immunotherapeutic agents. Here, a main advantage is that the model is on a B6 background, and it is compatible with many transplantable tumor cell lines, which will allow investigators to compare anti-tumor and irAE responses directly. The existence of irAE models will allows one to test the impacts of other relevant factors on the penetrance and severity of irAEs, such as genetic factors, diet, microbiome, etc. Finally, the NINJA model is modular, allowing investigators to study peripheral tolerance and irAEs in tissues of interest by utilizing animals with transgenic expression of Cre under different tissue-specific promoters. Ongoing work in our lab has focused on studying these processes in colon, pancreas, and liver, and, as discussed below, suggests that the outcome of tolerance will vary based on the tissue of self-Ag expression.

Previous studies have highlighted the role of ICRs, especially PD-1, in establishing peripheral tolerance through intrinsic regulation of self Ag-specific T cells that resulted in their functional unresponsiveness or deletion in experimental models of diabetes and encephalomyelitis (Keir *et al.*; Liang et al.; Martin-Orozco *et al.*; Sharpe et al.). The data presented in our study suggest that ICR-mediated regulation can also mediate immunological tolerance of functional self Ag-specific CD8 T cells within tissues. This does not preclude a potentially non-redundant role for ICR-mediated T cell regulation in the dLNs, as ligands for ICRs can be expressed by hematopoietic and non-hematopoietic cells both in peripheral organs and in their dLNs, thus potentially providing T cell regulation at multiple anatomical sites (Brown *et al.*; Diehl et al.; Keir *et al.*; Keir *et al.*; Latchman *et al.*; Liang *et al.*; Martin-Orozco *et al.*). In fact, recent evidence from our group and others shows that in cancer patients (Lucca et al.; Yost et al.) and tumor experimental models (Dammeijer et al.; Francis et al.; Kleinovink et al.; Connolly et al.) CPI-responsive tumor-specific CD8 T cells within tumor dLNs. Future studies will be required to understand whether migration of Ag-specific CD8 T cells from local dLNs is required for development of CPI-dependent tissue pathology in our model or whether Ag-specific CD8 T cells found in skin are sufficient to drive local tissue pathology following CPI administration.

One striking feature of skin Ag-specific CD8 T cells in our model was the similar gene expression signatures between cells from Ag-expressing mice with and without CPIs. This was true for most of the genes responsible for CD8 T cell effector functions (except *GzmA*) and strongly suggests that the T cells in these two conditions had similar effector capacity. This situation differs greatly from classical models of anergy/deletion, in which the expression of effector genes is not induced in the presence of Ag alone (Curtsinger et al.; Parish et al.; Redmond *et al.*). In N/C mice, skin Ag-specific CD8 T cells expressed high amounts of GrzB protein following Ag induction, and no significant change was observed upon CPI treatment. GrzB is a main cytolytic effector molecule for CD8 T cells and its high expression and production *in vivo* in our model confirms that the Ag-specific CD8 T cells in skin were not anergic. Furthermore, our data suggests that the ICR-mediated regulation of IFNγ and surface CD107a occurs after skin infiltration and at a post-transcriptional level, as protein expression was reduced in the skin compared to dLNs in Ag alone but not Ag + combo mice. Production of TNFα protein was observed in Ag + combo mice, however it was also correlated with the appearance of localized skin pathology at low penetrance in Ag only mice. This suggests that CPI treatment does not induce a unique functional state in self Ag-specific CD8 T cells, but rather it reduces their threshold for production of effector proteins that mediate tissue pathogenesis. This is not to say that CPI treatment did not alter the expression of any genes in Ag-specific CD8 T cells, as we did observe increases in *GzmA* mRNA and GrzA protein after CPI treatment, but the number of differentially upregulated genes in Ag + combo vs Ag alone mice was surprisingly low given the degree of tissue pathology observed. Interestingly, TNFα blockade in our model did not prevent CPI-induced skin pathology (data not shown), which is in sharp contrast with the beneficial effects of TNFα inhibitors observed in a model of DSS-induced and CPI-exacerbated colitis (Perez-Ruiz *et al.*). This difference suggests that different pathogenic mechanisms may drive irAEs depending on the affected organ.

Remodeling of local myeloid cells in tissues is observed in a variety of immunological challenges, including infections (Shi and Pamer), cancer (Ugel et al.), and in response to commensal bacteria (Bain and Mowat), and it is thought to be one of the compensatory mechanisms that tissues use to support and regulate an ongoing immune response (Nobs and Kopf). In our study, infiltration of effector-like Ag-specific CD8 T cells into Ag-expressing skin was also associated with a dramatic remodeling of the local myeloid compartment, yet the changes were almost imperceptible at the macroscopic and histological levels. The increase in frequency of skin CD11b+CD11c+ cells and their upregulation of ISGs, including *Cd274*, after local Ag induction indicate a crosstalk between tissue-infiltrating self Ag-specific effector-like CD8 T cells and myeloid cells is potentially occurring below the level of detection in healthy tissues. This is in line with recent CyTOF analysis of human fetal skin that revealed the presence of PD-L1+ myeloid cells and a small fraction of infiltrating CD45RO+ CD8 T cells that had higher expression of Nur77, Ki-67, and IFNγ (suggesting they could be self-reactive)(Dhariwala et al.). Future studies in humans and mice will be needed to gain a better understanding of how myeloid and parenchymal cells adapt to maintain T cell homeostasis in healthy tissues, and why this process breaks down in the context of autoimmunity and CPI therapy.

Because of its anatomical location and function, skin is considered a relatively immunologically active organ that is perhaps more prone towards driving CD8 T cell activation. Over the years, it has been debated whether skin-specific expression of Ag leads to CD8 T cell tolerance or activation, and many different experimental models of skin-specific Ag expression have been used for these studies. Skin is equipped with unique innate immune cell populations, like Langerhans cells (LCs) that patrol the epidermis, efficiently deliver Ag and inflammatory cues to skin dLNs, and express higher levels of co-stimulatory molecules than immature Ag-presenting cells found in lymphoid organs under steady-state conditions (Kashem et al.; Kissenpfennig et al.). Nonetheless, LC-specific expression and presentation of Ag has been associated with development of Ag-specific T cell anergy or deletion (Flacher et al.; Gomez de Agüero et al.; Igyarto et al.; Shklovskaya et al.; Strandt et al.), thus recapitulating the tolerogenic outcome of DC-dependent Ag presentation in the absence of inflammation. Likewise, many studies, but not all (Mayerova et al.; McGargill et al.), have shown that upon expression of model Ag in keratinocytes, Ag-specific CD8 T cells do not infiltrate the tissue or are deleted (Bianchi et al.; Holcmann et al.; Rosenblum et al.; Waithman et al.) but that this state of tolerance can be broken under strong inflammatory conditions, such as tissue damage, administration of mature Ag-loaded DCs, or depletion of Tregs. Our studies were not designed to test how self-Ag specific CD8 T cells are primed in our model, but the NINJA model provides a useful system for dissecting the contributions of different skin and/or myeloid cell types in this process.

Our findings in skin suggest that lichenoid irAEs occur because PD-1+ effector CD8 T cells in the skin target epithelial cells after PD-1 blockade and that this does not require genetic predisposition towards autoimmunity. irAEs occur in most CPI-treated patients, and in many locations, and with different CPIs causing different spectra of events. This would support a model where most healthy individuals have pathogenic effector CD8 T cells in their tissues, and that ICRs prevent these cells from causing pathology. This raises the question of why irAEs occur in some locations and not others. Unlike skin, experimental evidence that suggests that self-Ag specific CD8 T cells that migrate into the small intestine become anergic and that this state is not rescued by CPI treatment (Nelson *et al.*; Pauken *et al.*). Likewise, migration into pancreas has been associated with deletion, anergy, or ignorance, depending on the model (Parish et. al.). We have also observed CD8 T cell anergy in NINJA mice when self-Ag expression is induced in hepatocytes, and this is not reversed by acute LCMV infection with concurrent CPI treatment (M.D. and N.S.J., manuscript in preparation). This suggests that peripheral tolerance mechanisms may vary based on the cell type or organs expressing the self Ag, with some encounters inducing more traditional T cell-intrinsic tolerance mechanisms and others relying on T cell-extrinsic mechanisms to prevent immunopathology. In the latter setting, ICRs will likely be important for establishing and maintaining tolerance, with potentially different ICRs being important in different tissues. Together, this model could help explain the spectrum of irAEs observed between patients and between antibodies targeting the PD-1/PD-L1 and CTLA4 pathways.

## Supporting information

All supplemental figs

## Acknowledgements

We thank Joshi lab members and D. Schatz, M. Bosenberg, Ian Parish, and Stephanie Eisenbarth for helpful discussions and reviewing the manuscript. We also thank the Yale Flow Cytometry Core, Yale School of Medicine Comparative Pathology Research Core, the Yale Center for Genome Analysis, and M. Huelsmeyer for tool supply. This work was supported by grants from The G. Harold & Leila Y. Mathers Foundation (YAL182 to N.S.J.), the Melanoma Research Alliance (569588 to N.S.J.) and the Yale Skin Cancer Spore-CEP (2P50CA196530-06 to N.S.J.). M.D. is supported by the Leslie H. Warner Post-Doctoral Research Fellowship.

## Author contributions

M.D. and N.S.J. designed research and wrote paper. M.D. analyzed data. M.D., I.W., C.C., D.K., N.I.H., and K.C. performed research, W.E.D. and J.S.L. provided histology data from human samples. All authors have read and approved the manuscript.

---

**Figure S1 - Examples of skin pathology and matching to clinical severity scoring in experimental mice.** Representative pictures and correspondent clinical severity score assigned to mice across experimental groups.

**Figure S2 - EpCAM+CD45− epidermal cells constitute most of Ag-expressing skin cells and are subjected to CD8 T cell-mediated killing after CPI treatment.**

A) Flow cytometric analysis of total GFP+ (Ag-expressing) skin cells from the indicated experimental groups and analyzed by EpCAM and CD45 expression. Numbers represent average ± standard deviation of gated populations (EpCAM+CD45− = epidermal keratinocytes; CD45+EpCAM+ = Langerhans cells; CD45+EpCAM− = all other hematopoietic skin cells). B) Total numbers of GFP+ cells harvested from the skin of mice treated as indicated. For comparisons of untreated vs + Ag groups, ** *P* = 0.0019 (CD45+EpCAM-), *** *P* = 0.0009 (CD45+EpCAM+) and **** *P* < 0.0001 (EpCAM+CD45-) by *t* test. For comparisons of + Ag vs + Ag + combo groups, ** *P* = 0.0018 (CD45+EpCAM+), **** *P* < 0.0001 (EpCAM+CD45-) and ns = not significant (CD45+EpCAM-) by *t* test. For comparisons of + Ag + combo vs + Ag + combo + αCD8 groups, **** *P* < 0.0001 (EpCAM+CD45-) and ns = not significant (CD45+EpCAM- and CD45+EpCAM+) by *t* test. n = 5-7, pool of 2 experimental repeats, representative of 3 independent repeats.

**Figure S3 - Localized lichenoid immuno-mediated pathology develops in mice adoptively transferred with transgenic Ag-specific CD8 T cells (P14 CD8 T cells) following skin-restricted Ag expression and systemic CPI administration.**

A) Experimental schedule for adoptive transfer of P14 CD8 T cells into N/C mice followed by Ag induction and CPI treatment. B) Representative pictures of macroscopic skin pathology in mice adoptively transferred with P14 CD8 T cells and treated as indicated. C) Distribution of clinical severity scores for the skin of P14 CD8 T cell-transferred mice from the indicated experimental groups. Median shown in dot plot. * *P* = 0.0179 (untreated vs combo CPIs) or 0.0261 (untreated vs + Ag), and **** *P* < 0.0001 by *t* test. n = 6-14, pool of 2 experimental repeats, representative of > 3 independent repeats.

**Figure S4 - Mediators of T cell effector functions are produced by endogenous Ag-specific CD8 T cells following skin-specific Ag induction.**

Example flow cytometric analysis of endogenous GP33-specific CD8 T cells harvested from the dLNs and *ex vivo* peptide stimulated (top) or skin without ex vivo stimulation (bottom) of N/C mice treated as indicated. Numbers in plots represent average ± standard deviation of frequency of the corresponding gated population. n = 5, representative of > 3 independent repeats.

**Fig S5 - Skin-infiltrating Ag-specific CD8 T cells transcriptionally express several ICRs after localized Ag induction.**

ICR gene expression heatmap for P14 CD8 T cells and total CD8 T cells harvested from Tam-treated skin of mice from the indicated experimental conditions and analyzed by scRNAseq. n = 884 cells (total skin CD8 T cells from “No Ag no CPIs” group), 1203 cells (P14 CD8 T cells from “+ Ag no CPIs” group), 2053 cells (P14 CD8 T cells from “+ Ag + combo” group), 2201 cells (total skin CD8 T cells from “+ Ag no CPIs” group), 2565 cells (total skin CD8 T cells from “+ Ag + combo” group).

**Figure S6 - Phenotypic and CD8 T cell functional differences between scRNAseq experimental groups reflect the expected immunological outcome based on treatment.**

Skin appearance and clinical severity score are shown for each experimental mouse included in the scRNAseq analysis from Figures 3-4. Total skin cells harvested from the area highlighted by the red rectangle (Tam-treated) from each mouse were analyzed for presence of P14 CD8 T cells (dsRED+) and Ag (GP33)-specific functional responses prior to sequencing. Numbers in facs plots indicate frequency of the corresponding gated population, n = 2.

**Fig S7 - Skin localized tissue pathology develops with low penetrance after induction of Ag in the absence of CPI administration.**

Example skin appearance of N/C mice treated with Dox-containing food and Tam applied to the skin area indicated by the red rectangle. Frequency and clinical severity score for each skin phenotype are shown with corresponding functional data on Ag-specific endogenous (top) or transgenic (bottom) CD8 T cells. Numbers in facs plots represent average ± standard deviation of frequency of the corresponding gated population. n > 20, pool of > 3 independent repeats.

**Figure S8 - *CD274*/PD-L1 is not expressed by EpCAM+CD45− skin cells.**

A) Expression of *CD274* (PD-L1) across all skin cells harvested from the indicated experimental conditions and clustered as shown in Fig. 3B following scRNAseq analysis. n = 10162 cells (total skin cells from “No Ag no CPIs” group), 9017 cells (total skin cells from “+ Ag no CPIs” group), 6019 cells (total skin cells from “+ Ag + combo” group). B) Representative flow cytometric analysis of total skin cells harvested from N/C mice treated as indicated and analyzed for EpCAM, CD45 and PD-L1 expression. Numbers in EpCAM/CD45 plot indicate average ± standard deviation of frequency of EpCAM+CD45− total cells from Tam-treated skin. Numbers in histogram indicate average ± standard deviation of PD-L1 MFI by EpCAM+CD45− skin cells from the conditions analyzed. n = 5, representative of > 3 independent repeats. C) Representative flow cytometric analysis of total skin cells harvested from mice treated as indicated and analyzed for GFP (Ag) and PD-L1 expression. Numbers in SSC/GFP plot indicate average ± standard deviation of frequency of GFP+ total cells from Tam-treated skin. Numbers in histogram indicate average ± standard deviation of PD-L1 MFI by GFP+ skin cells from the conditions analyzed. n = 5, representative of > 3 independent repeats. D) Hallmark gene set score for IFN-stimulated genes (“ISG score”) computed for all skin cells harvested from the indicated experimental conditions and clustered as shown in Fig. 3B following scRNAseq analysis. n = 10162 cells (total skin cells from “No Ag no CPIs” group), 9017 cells (total skin cells from “+ Ag no CPIs” group), 6019 cells (total skin cells from “+ Ag + combo” group).

**Figure S9 - Low expression of *Ifngr1* and *Ifngr2* is detected in skin epithelial cells following localized Ag induction.**

Expression of *Ifngr1* (top) and *Ifngr2* (bottom) across all skin cells harvested from the indicated experimental conditions and clustered as shown in Fig. 3B following scRNAseq analysis. n = 10162 cells (total skin cells from “No Ag no CPIs” group), 9017 cells (total skin cells from “+ Ag no CPIs” group), 6019 cells (total skin cells from “+ Ag + combo” group).

**Figure S10 - Sub-clustering of skin myeloid cells.**

Bubble plot of canonical marker genes utilized to identify sub-clusters of skin myeloid cells analyzed as described in Fig. 4A-B.

**Figure S11 – Additional transcriptomic analysis of P14 CD8 T cells.**

A) Pie charts shown the fraction of *dsRED*-expressing cells (P14 CD8 T cells) in the indicated clusters n = 1203 (P14 CD8 T cells from “+ Ag no CPIs” group) and 2053 (P14 CD8 T cells from “+ Ag + combo” group). B) Scatterplots of average gene expression in P14 CD8 T cells in the indicated clusters from “+ Ag no CPIs” (x) and “+ Ag + combo” (y) experimental conditions. C) Distribution of CD8 T cell effector scores calculated as in Figure 3I, but omitting *Gzma*. **** *P* < 0.0001 by Wilcoxon rank sum test.

**Table S1 - List of primers.**

## Methods

### Mouse studies

All studies were carried out in accordance with the guidelines of the Declaration of Helsinki and approved by the Institutional Animal Care and Use Committees of Yale University (IACUC 2019_20112). All mice were bred in specific pathogen-free conditions. For experiments, 6-12 week old female and male mice were used.

### Human samples

Cases of lichenoid cutaneous irAEs were identified by searching the Yale Dermatopathology clinical case database. Selected cases were de-identified and H&E-stained slides from clinical biopsy samples were imaged. These studies were conducted according to the guidelines of the Declaration of Helsinki and approved by the Yale Institutional Review Board (IRB protocol ID 1501015235, approved on 2/26/2021).

### Generation and genotyping of NINJA strains, genotyping of TCR transgenic P14 mice

NINJA mice were generated as previously described (Damo *et al.*). NINJA x CAG-rtTA3 were generated by crossing NINJA mice to CAG-rtTA3 mice (JAX, stock # 016532) and are henceforth referred to as NINJA. N/C (*Rosa26-NINJA/CreER^T2^;CAG-rtTA3 Tg*) mice were generated by crossing NINJA mice to R26-CreER^T2^ mice (JAX, stock # 008463). Thy1.1/1.1+ fLuciferase- or dsRED-expressing TCR transgenic P14 mice (Pircher *et al.*) were bred in-house.

For genotyping, mouse tail DNA was amplified using GoTaq G2 Green Master Mix (Promega, cat. M7823) or KAPA Taq PCR Kit (Takara Bio, cat. R040A). Primers Rosa26 WT F, Rosa26 WT R, and Rosa26 Ninja R (**Table S1**) were used for genotyping of the NINJA allele, with expected bands of 378bp for WT allele and 280 bp for NINJA allele. For CAG-rtTA3 genotyping, primers CAGrtTA common F, CAGrtTA WT R and CAGrtTA Transgene R (**Table S1**) were used and expected bands were 360bp for WT allele and 330bp for Transgene. For R26-CreER^T2^ genotyping, primers generic-Cre_1, generic-Cre_2, generic-Cre_3 and generic-Cre_4 (**Table S1**) were used with expected bands of 500bp for internal control and 320bp for Transgene. For P14 genotyping, primers P14r F and P14r R (**Table S1**) were used with expected band of 300 bp for Transgene. For fLuciferase genotyping, primers 10946, 10947, oIMR7338 and oIMR7339 (**Table S1**) were used with expected bands of 324bp for internal control and 200bp for Transgene. For dsRED genotyping, primers oIMR3847, oIMR4110, oIMR7338 and oIMR7339 (**Table S1**) were used and expected bands were 324bp for internal control and 208bp for Transgene.

### In vivo *induction of NINJA antigens in the skin and antibody treatments*

Mice were fed doxycycline-containing food (Envigo Teklad, cat. TD.120769) for ten consecutive days. Mice also received four doses of a 50mg/mL solution of 4-hydroxy-tamoxifen (Millipore Sigma, cat. T176) in DMSO (AmericanBIO, cat. AB03091) that was painted on the indicated area of the skin on the lower back of each experimental mouse (10-20uL/dose) every other day starting the day after addition of doxycycline diet.

For CPI treatment, PD-1 blocking antibody (clone 29F.1A12, BioXCell, cat. BP0273) and CTLA-4 blocking antibody (clone 9D9, BioXCell, cat. BE0164) were administered either in combination or as single agents at 100ug/antibody/mouse every third day starting on the day of doxycycline diet addition for four total doses.

For T cell depletion, CD8 depleting antibody (clone 53-6.7, BioXCell, cat. BP0004-1) and CD4 depleting antibody (clone GK1.5, BioXCell, cat. BP0003-1) were administered either in combination or as single agents at 200ug/antibody/mouse every third day starting one week prior to doxycycline diet addition for seven total doses.

For IFNγ-specific blockade, IFNγ blocking antibody (clone XMG1.2, BioXCell, cat. BE0055) was administered at 250ug/mouse every third day starting on the day of doxycycline diet addition for six total doses.

### *Preparation of skin and LN cells*, in vitro *antigen-specific restimulation and flow cytometry*

4-hydroxy-tamoxifen-treated skin was harvested at experimental endpoint and processed using the mouse epidermis dissociation kit according to manufacturer’s instructions (Miltenyi, cat. 130-095-928). LN cells were harvested and processed as previously described (Damo *et al.*). For antigen-specific restimulation, 1 * 10^6^ cells from skin or LNs were cultured for 6hr at 37°C in 5% CO_2_ in complete RPMI-1640 (10% HI-FBS, 55μM beta-mercaptoethanol, 1x Pen/Strep and 1x L-Glut) supplemented with LCMV GP33-41 peptide (0.5μg/mL, AnaSpec, cat. AS-61296) or left unstimulated. For restimulation of skin lymphocytes, 10^4^ wild-type congenically-marked donor LN APCs were added to the co-culture.

For flow cytometric analysis, 1-5 * 10^6^ cells were stained with antibodies specific for surface markers Thy1.2 (clone 30-H12, BioLegend), Thy1.1 (clone OX-7, BioLegend), CD8α (clone 53-6.7, BioLegend), CD8β (clone 53-5.8, BioLegend), CD4 (clone GK1.5 or RM4-4, BioLegend), CD44 (clone IM7, BioLegend), tetramers for H2Db/GP33-41-specific CD8 T cells (from NIH Tetramer Core Facility), PD-1 (clone 29F.1A12, BioLegend), CTLA-4 (clone UC10-4B9, BioLegend), EpCAM (clone G8.8, ThermoFisher Scientific), CD45 (clone 30-F11, BioLegend), CD11b (clone M1/70, BioLegend), CD11c (clone N418, BioLegend), PD-L1 (clone 10F.9G2, BioLegend), and CD107a (clone 1D4B, BioLegend). For intracellular staining, cells were processed using the Fixation/Permeabilization kit from BD Biosciences (BD Cytofix/Cytoperm, cat. 554714) following manufacturer’s instructions and stained with antibodies specific for GrzA (clone 3G8.5, BioLegend), GrzB (clone GB11, ThermoFisher Scientific), IFNγ (clone XMG1.2, BD Biosciences) and TNFα (clone MP6-XT22, eBioscience). Samples were analysed on a BD Biosciences LSRII flow cytometer.

### In vivo *imaging by IVIS*

For *in vivo* imaging of mice following skin-specific antigen induction, hair was removed from back of mice and mice were adoptively transferred with 5 x 10^3^ fLuciferase^+^ P14 T cells by retro-orbital injection (I.V.) one day prior to initiating doxycycline/4-hydroxy-tamoxifen regimen. Two, five, ten or fourteen days later mice were injected I.V. with a PBS solution containing 3mg of luciferin/mouse (XenoLight D-Luciferin - K+ Salt, PerkinElmer, cat. 122799) and imaged using an *In Vivo* Imaging system (IVIS, PerkinElmer) instrument.

### IF/IHC

For IF, mice were adoptively transferred with 5 x 10^3^ dsRED^+^ P14 T cells by retro-orbital injection (I.V.) one day prior to initiating doxycycline/4-hydroxy-tamoxifen regimen. Mice were euthanized at experimental endpoint to collect 4-hydroxytamoxifen-treated skin. Tissue was fixed overnight at 4°C with a paraformaldehyde-lysine-periodate fixative (PLP), subsequently cryoprotected with 30% sucrose in PBS for 6-8hr at 4°C, and then embedded in cryomolds with 100% optimum cutting temperature (O.C.T.) compound (VWR, cat. 25608-930) on dry ice for freezing. Tissue sections were processed with Vectashield Antifade mounting medium with DAPI (Vector Laboratories, cat. H-1200-10) and imaged on a Leica confocal microscope.

For IHC, skin tissues were fixed in a 1x formaldehyde solution in PBS (Millipore-Sigma, cat. 252549) and then embedded in paraffin and sectioned by the Yale School of Medicine Histology Core. All skin tissues were sectioned and stained (H&E or CD3) as previously described (Damo *et al.*). Imaging of IHC samples was performed using an EVOS Imaging System (ThermoFisher Scientific).

### Epidermal thickness score calculation

H&E-stained slides were obtained and imaged from each experimental sample as described above. Three independent ROIs were considered in each H&E image from various locations along the epidermis. Epidermal thickness was then calculated within each ROI by The Histological Inflammation Computation (THIC) analysis in MATLAB (version R2020b, 9.9.0.1467703).

Briefly, the pixel area and perimeter of each ROI were defined by the total number of white pixels and the boundary length of the observed binary epidermis. Thickness score for each ROI was then calculated as a ratio of pixel area by the perimeter length.

MATLAB code for initializing the THIC GUI and project files are available to use for research purposes only at https://github.com/dakwok/The-Histological-Inflammation-Computation-software.

### Cutaneous clinical severity scoring calculation

The scoring method is based on the Lichen Planus Severity Index (Kaur et al.). Briefly, each clinical severity score is calculated at experimental endpoint by assigning weights to the various morphologies of skin pathology: erythema (weight = 1), scale (weight = 1), lichenification (weight = 2), and erosion/ulceration (weight = 4). For each morphology seen in a given experimental mouse, the weight of that morphology is multiplied by the percentage of “at-risk” skin involved, defined as the area treated with topical Tam. These products are calculated for each morphology of lesion, and the results are summed, giving the final clinical severity score.

### Single-cell RNA sequencing

Total skin cells were prepared as described above. Single cell RNA sequencing libraries were prepared using Chromium Single Cell 5’ Reagent Kit (10X Genomics), according to manufacturer’s instructions. Sequencing was performed on an Illumina NovaSeq 6000 system. Data were processed using the 10X Genomics Cell Ranger (version 4.0.0) pipeline. For each sample, the gene-barcode matrices were passed to R (version 4.0.3) software package Seurat (version 4.0.1). Number of genes, UMIs, and the percentage of mitochondrial genes were plotted, and outliers were removed to filter out doublets and dead cells. 25,198 mouse single cells from two experimental repeats were pooled for further analyses (10,162 cells from “No Ag no CPIs” condition, 9,017 cells from “+ Ag no CPIs” condition and 6,019 cells from “+ Ag + combo” condition). Raw gene counts were log-normalized by Seurat function *NormalizeData* with parameter *normalization.method* set to “LogNormalize”. For the heatmaps of gene expression, relative abundances were calculated with Z-scaled average of log-normalized expression. Effector score for CD8 T cells was calculated by applying Seurat function *AddModuleScore* to the following gene list: *Gzma, Gzmb, Gzmk, Fasl, Prf1, Ifng, Tnf*. ISG score was calculated by applying *AddModuleScore* to the following gene list: *Stat1, Irf1, Cd274, Fcgr1, Ccr5, Cxcl9, Cxcl10, Gbp2, Gbp3, Gbp5, Ifitm1, Ifit1, Ifit2, Ifit3* (Ayers *et al.*; Benci *et al.*; Combes *et al.*; Liu *et al.*; Luoma *et al.*; Szabo *et al.*).

### Quantification, statistical analysis and renderings

FlowJo v10.7.1 software was used for flow cytometric analysis. Prism v.9.1.0 software was used to determine statistical significance using Student’s two-tailed unpaired *t* test, Wilcoxon rank sum test with Bonferroni correction or simple linear regression analysis. *P* values are indicated in each figure legend. BioRender was used for Fig 1B, Fig 3A and Fig S3.

