## Supplementary material for "The PD-1 checkpoint receptor maintains tolerance of self-reactive CD8 T cell in skin": All supplemental figs

**Figure S1: Examples of skin pathology and matching to clinical severity scoring in experimental mice.**

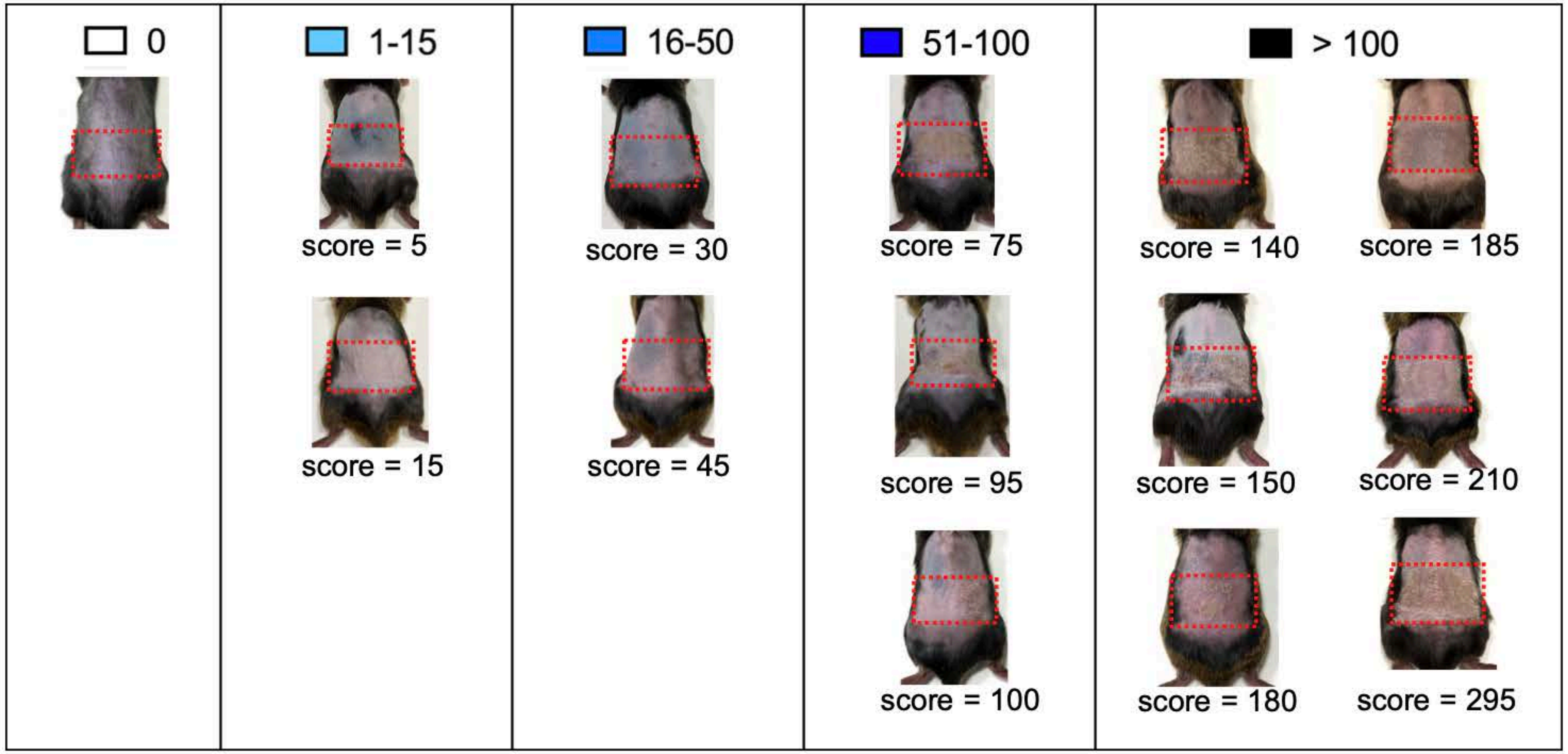

**Fig S2: EpCAM+CD45- epidermal cells constitute most of Ag-expressing skin cells and are subjected to CD8 T cell-mediated killing after CPI treatment.**

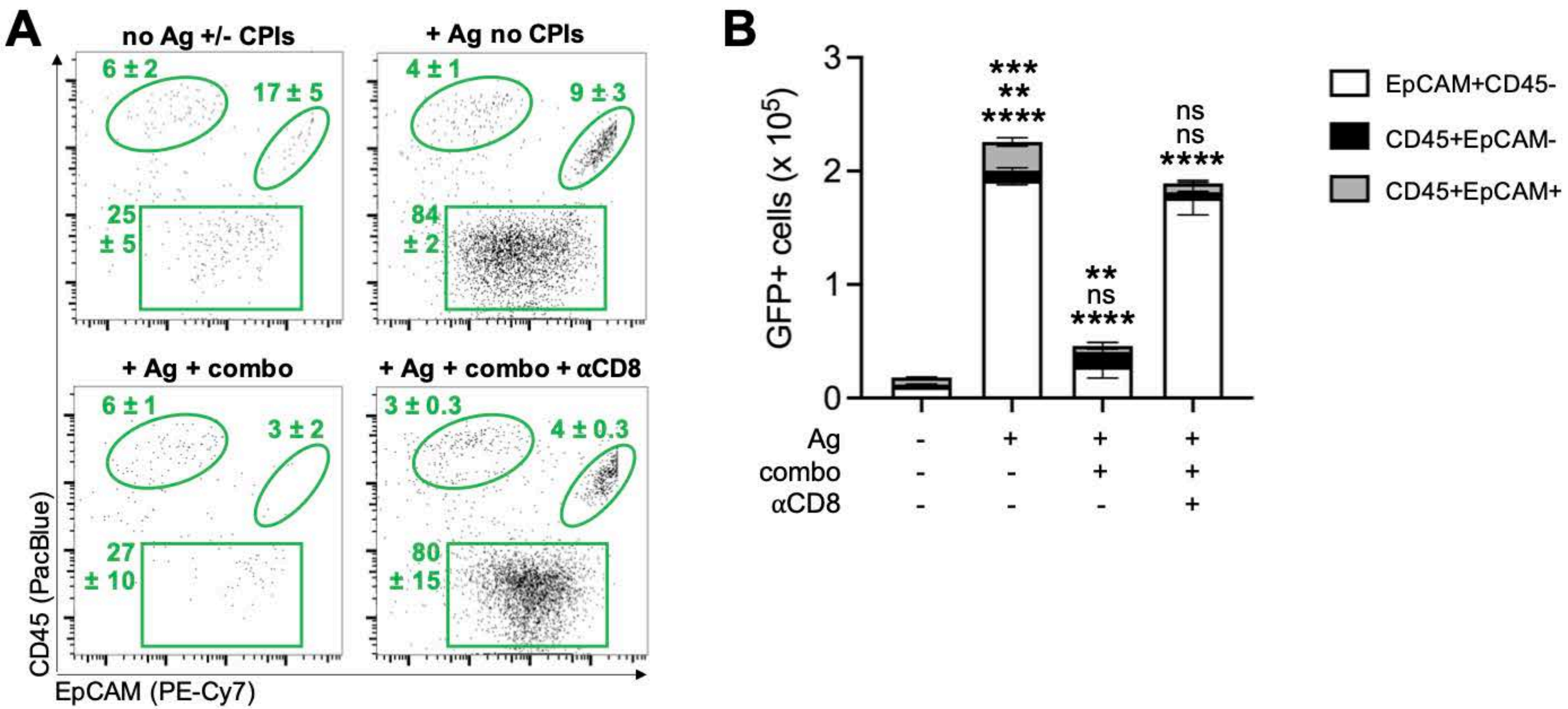

**Figure S3: Localized lichenoid immuno-mediated pathology develops in mice adoptively transferred with transgenic Ag-specific CD8 T cells (P14 CD8 T cells) following skin-restricted Ag expression and systemic CPI administration.**

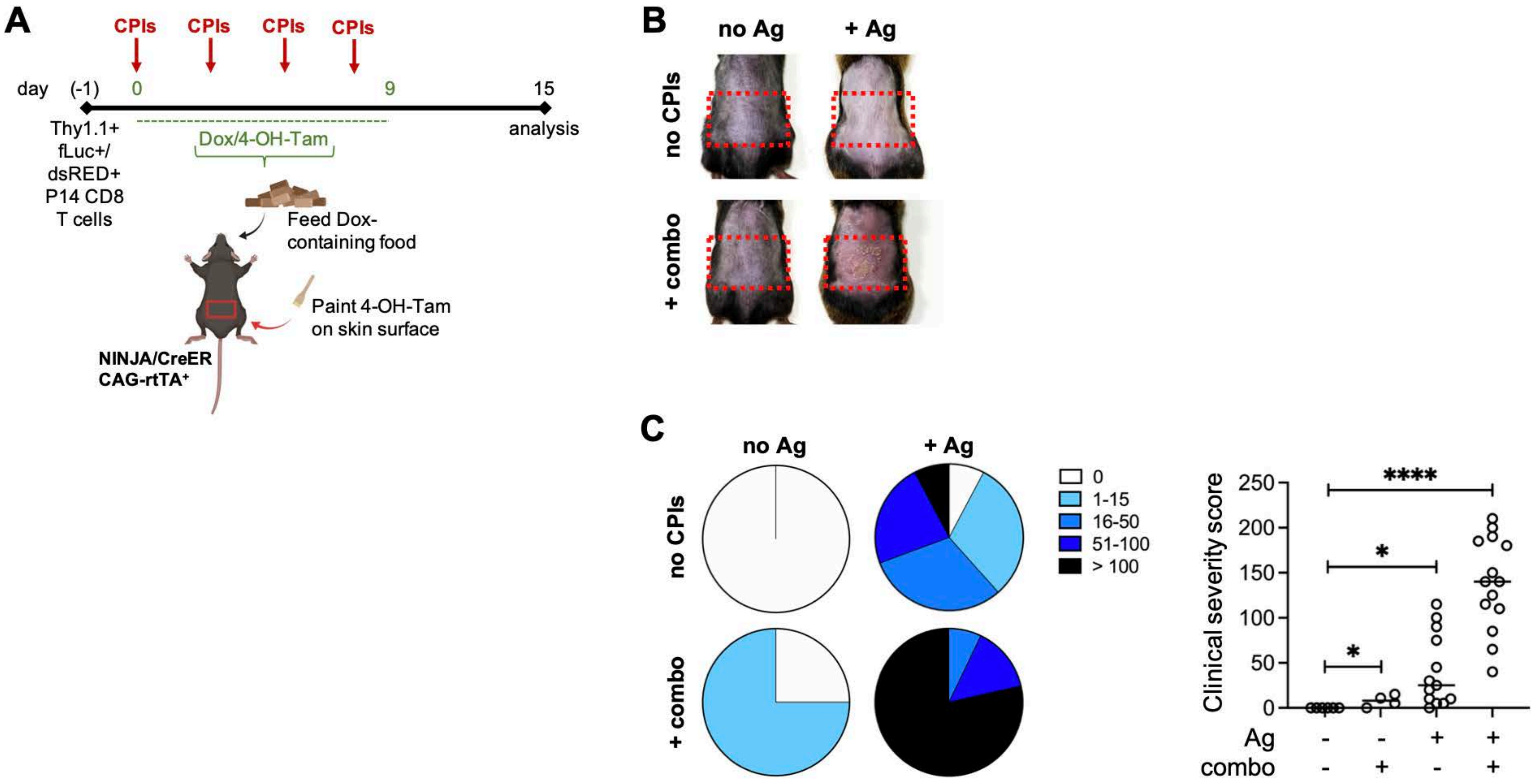

**Figure S4: Mediators of T cell effector functions are produced by endogenous Ag-specific CD8 T cells following skin-specific Ag induction.**

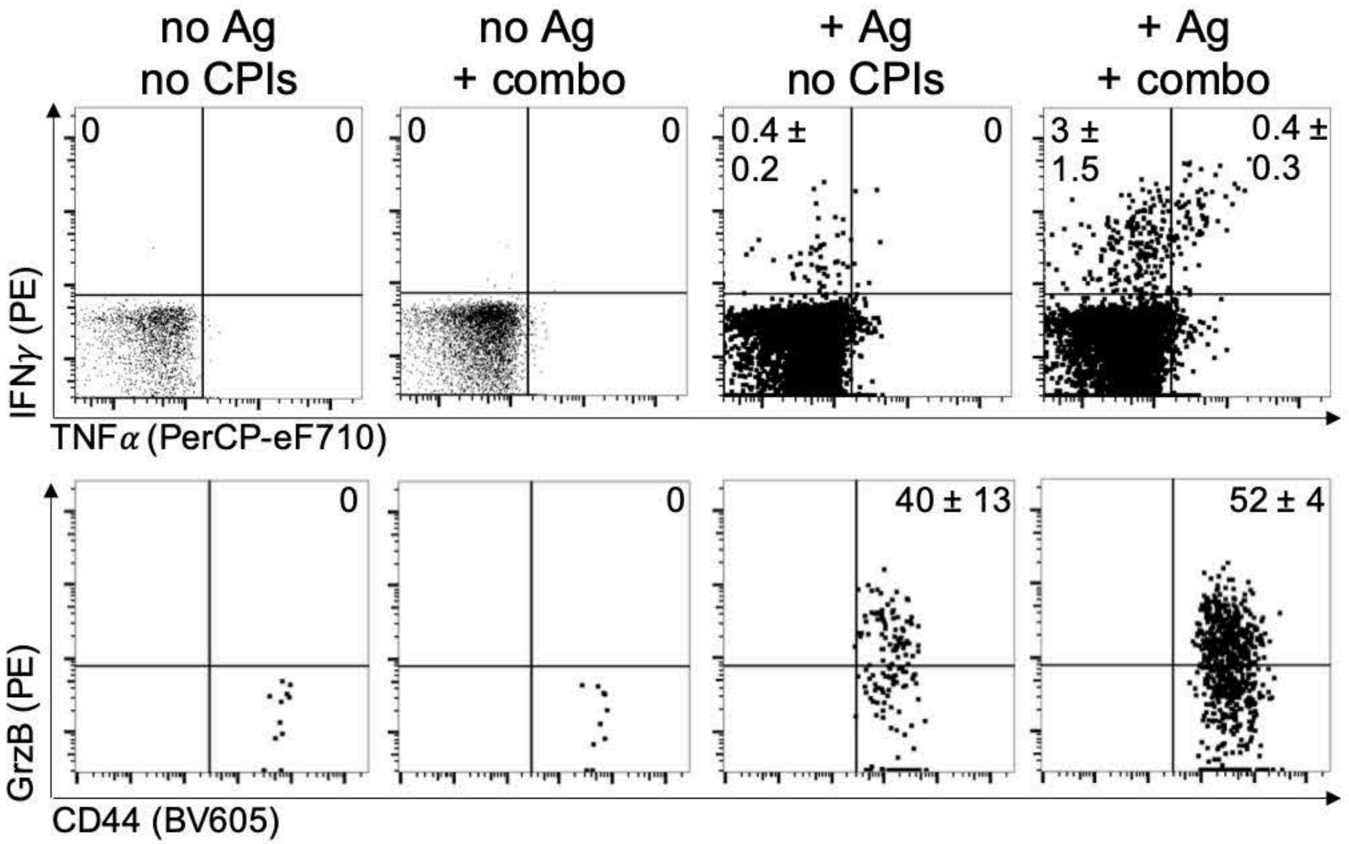

### Figure S5: Skin-infiltrating Ag-specific CD8 T cells transcriptionally express several ICRs after localized Ag induction.

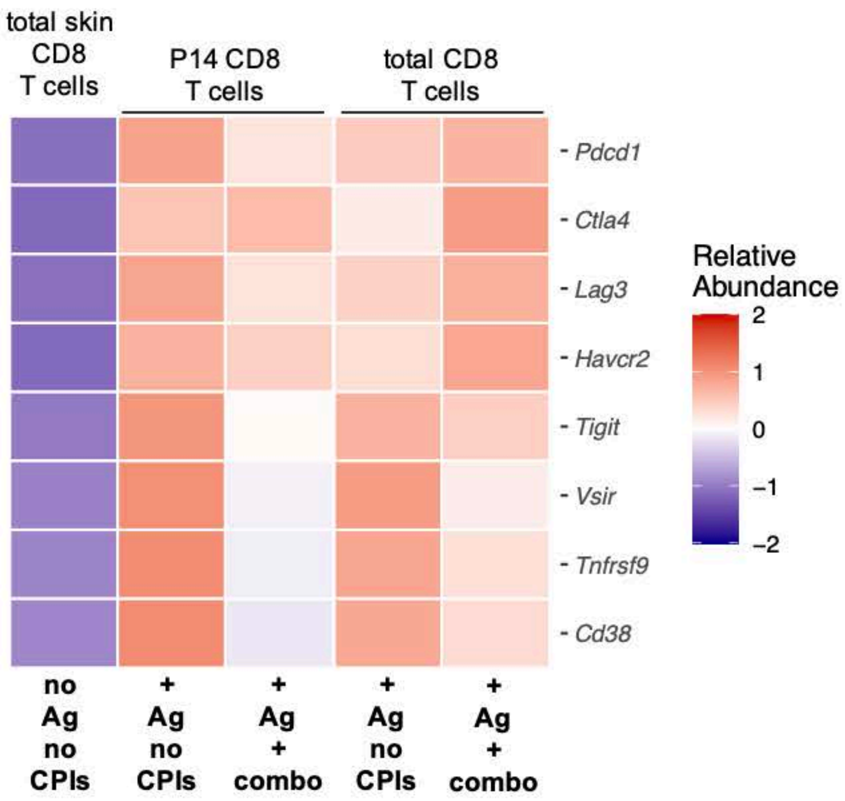

**Figure S6: Phenotypic and CD8 T cell functional differences between scRNAseq experimental groups reflect the expected immunological outcome based on treatment.**

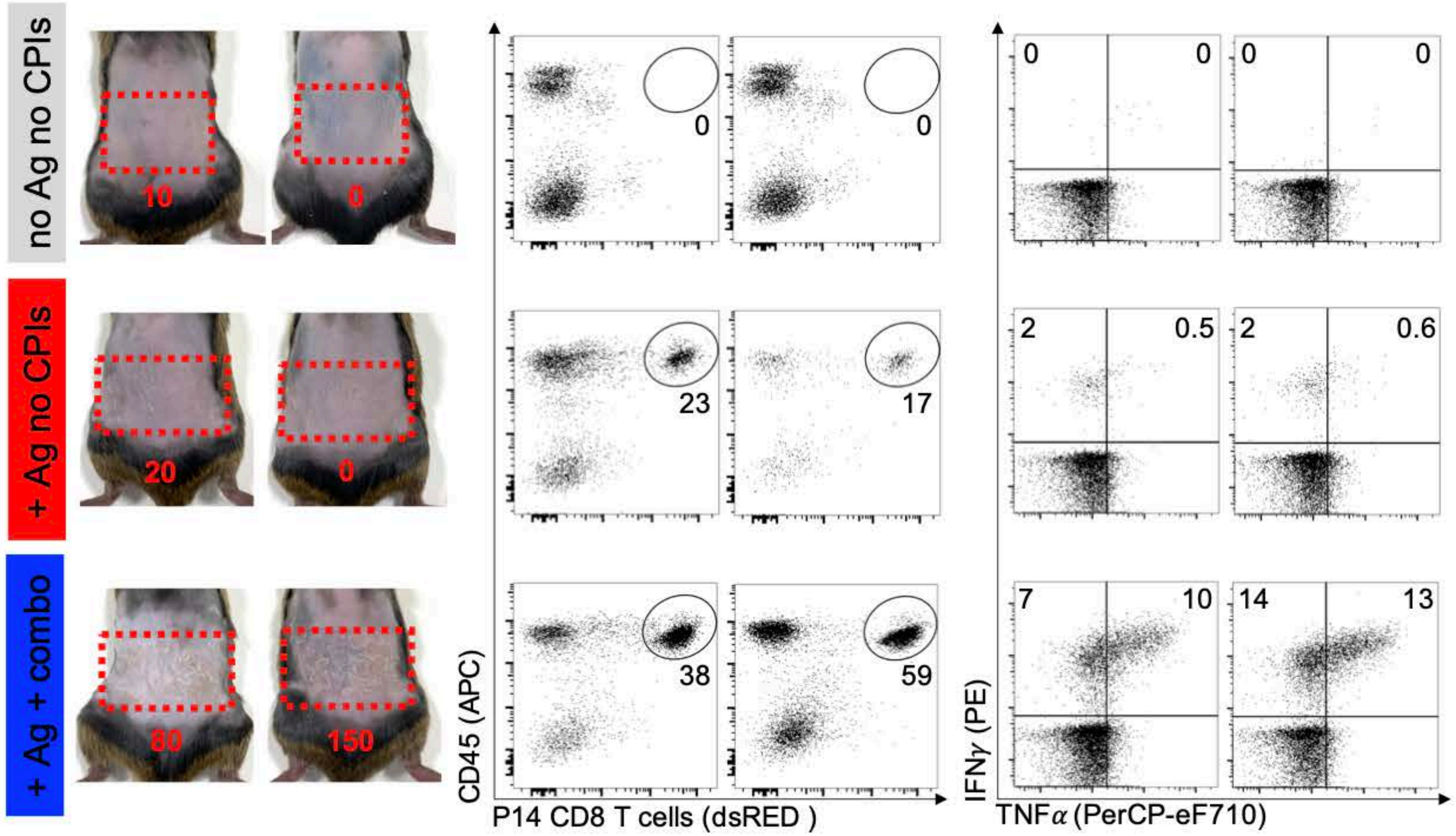

**Figure S7: Skin localized tissue pathology develops with low penetrance after induction of Ag in the absence of systemic CPI administration.**

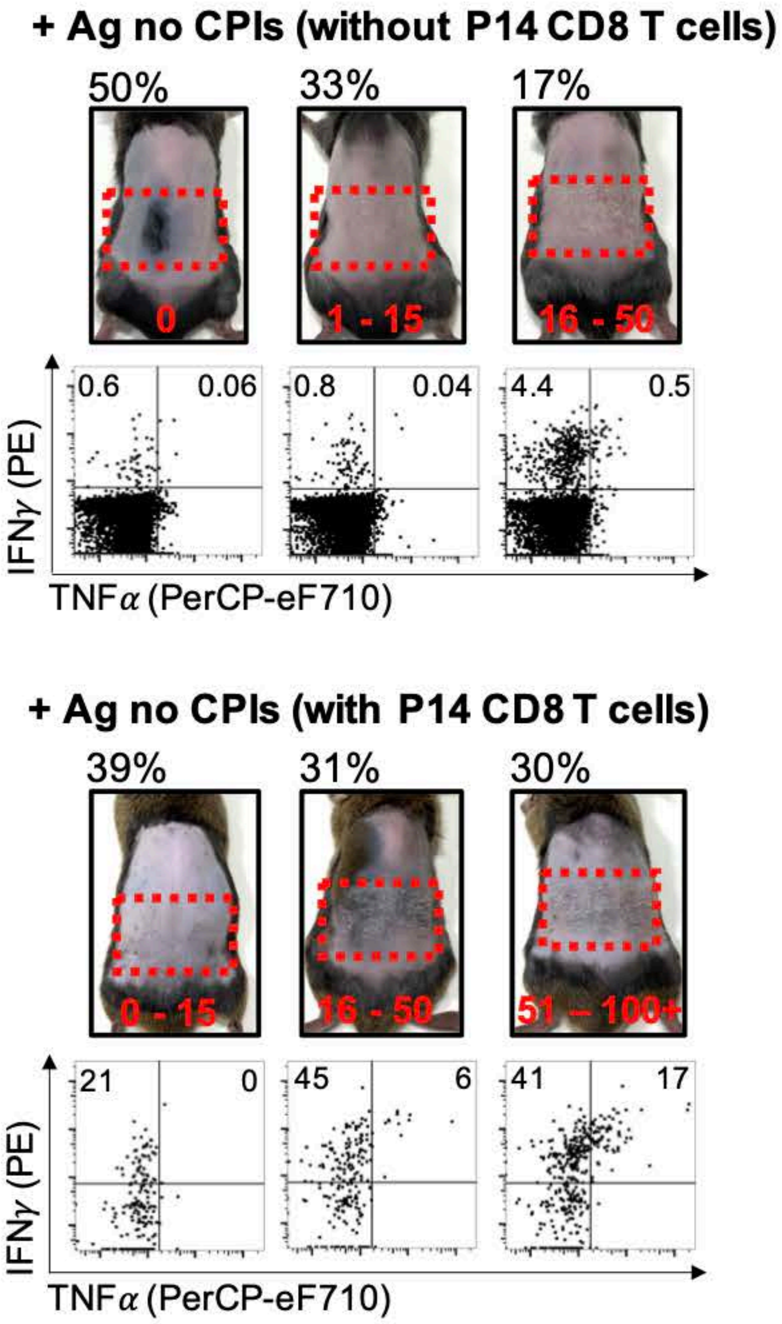

**Fig S8: *CD274*/PD-L1 is not expressed by EpCAM+CD45- skin cells.**

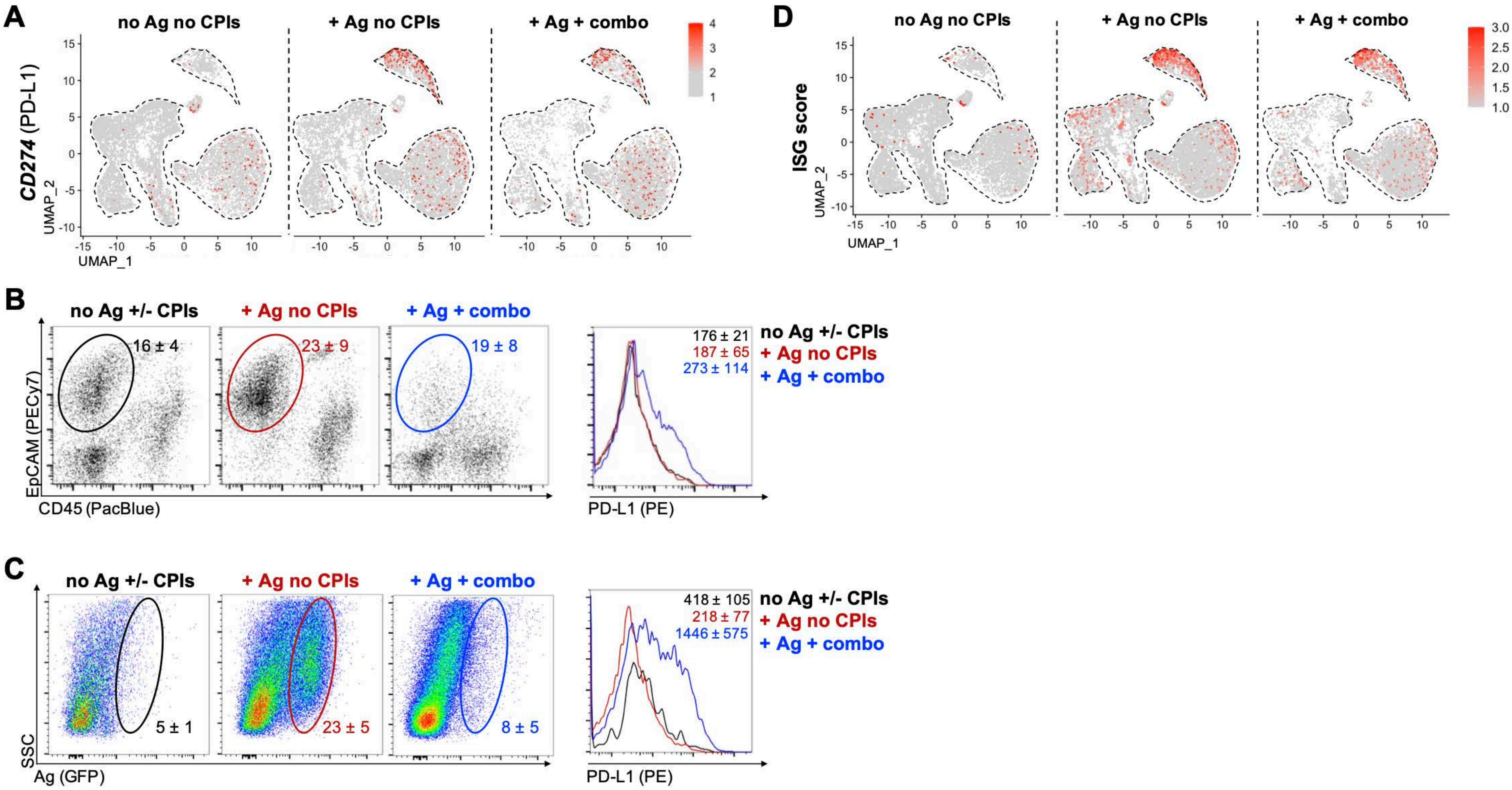

**Fig S9: Low expression of *Ifngr1* and *Ifngr2* is detected in skin epithelial cells following localized Ag induction.**

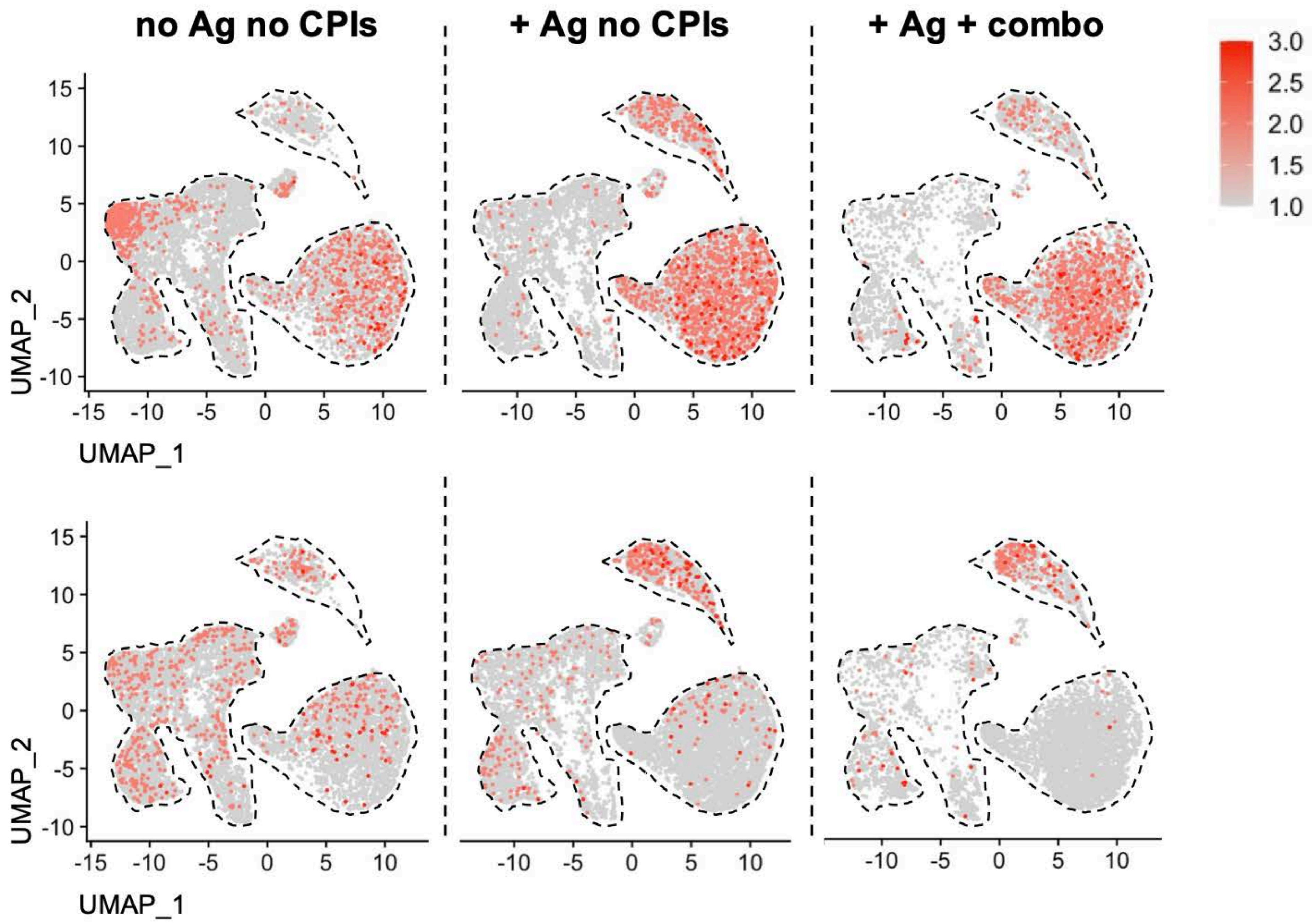

**Fig S10: Sub-clustering of skin myeloid cells.**

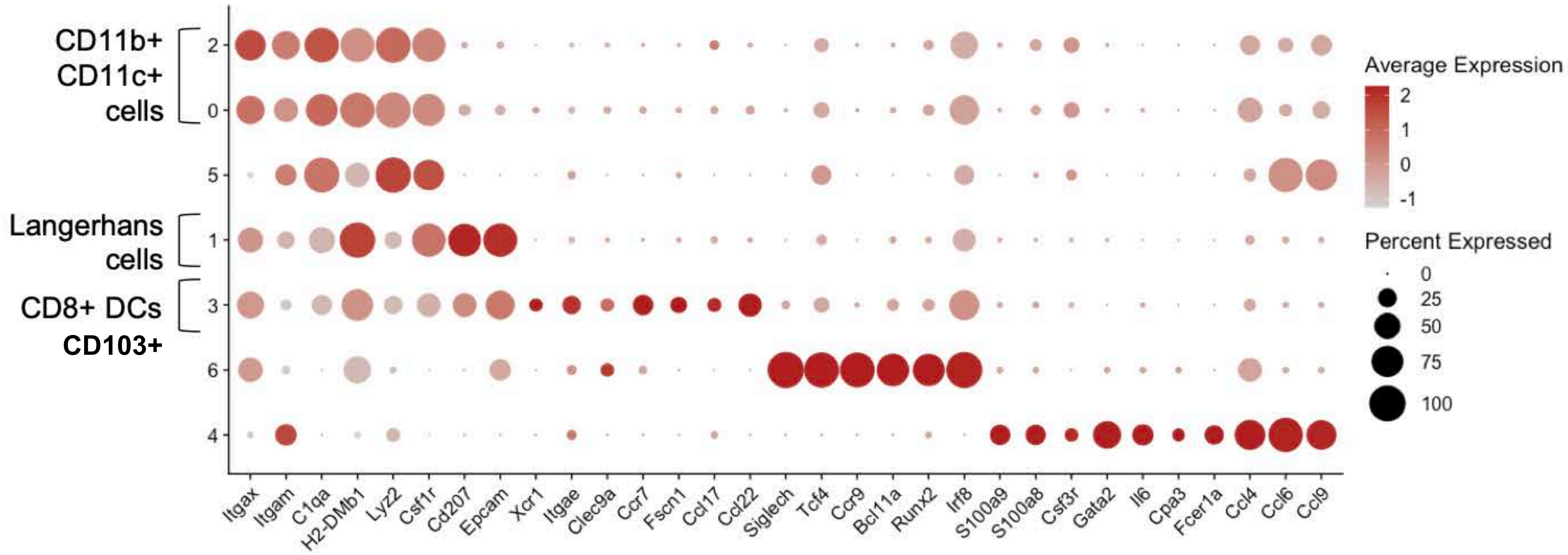

**Fig S11: Supplemental analysis of P14 CD8 T cells**

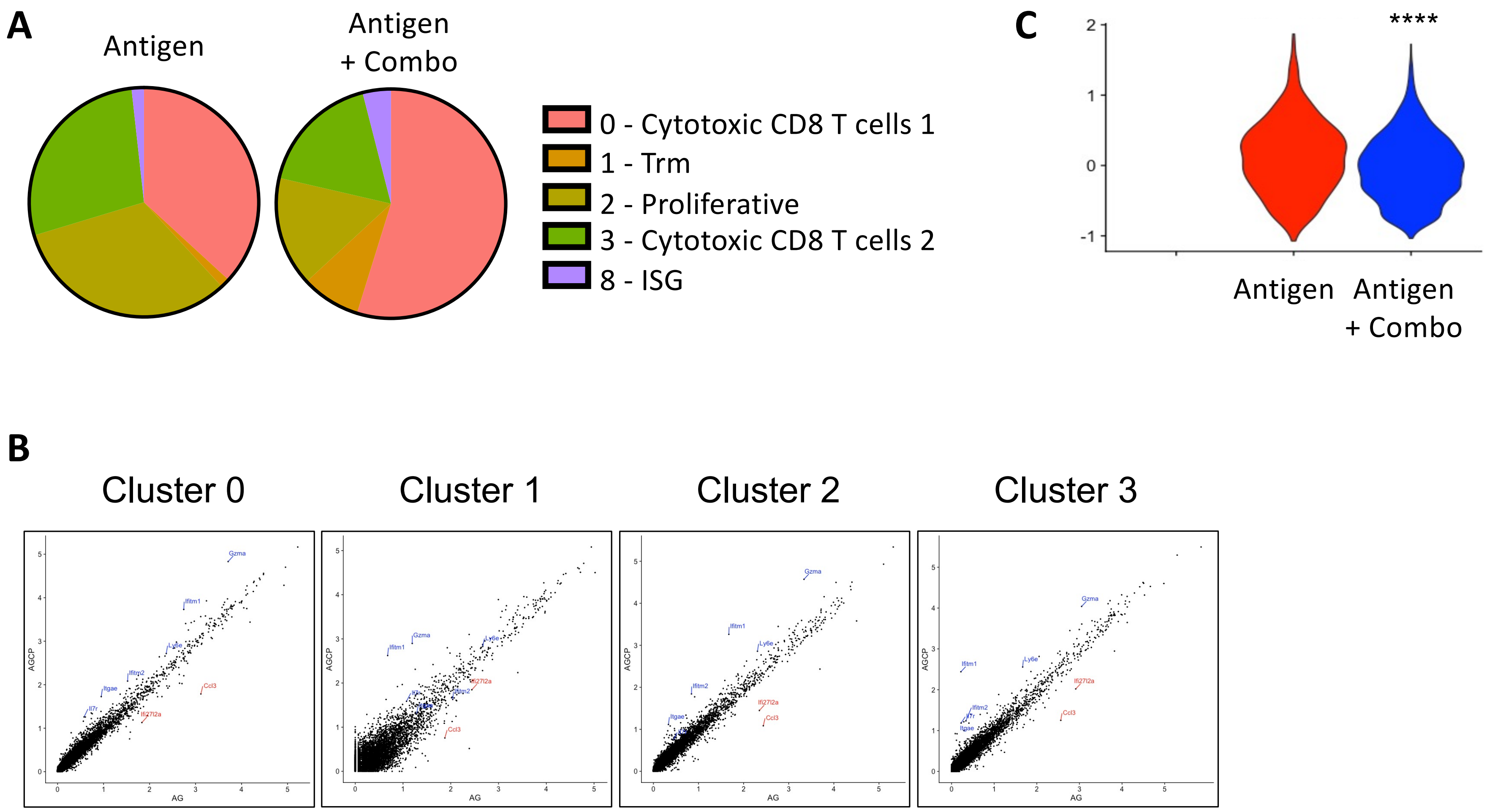
